# Soil variation response is mediated by growth trajectories rather than functional traits in a widespread pioneer Neotropical tree

**DOI:** 10.1101/351197

**Authors:** Sébastien Levionnois, Niklas Tysklind, Eric Nicolini, Bruno Ferry, Valérie Troispoux, Gilles Le Moguedec, Hélène Morel, Clément Stahl, Sabrina Coste, Henri Caron, Patrick Heuret

## Abstract

1. Trait-environment relationships have been described at the community level across tree species. However, whether interspecific trait-environment relationships are consistent at the intraspecific level is yet unknown. Moreover, we do not know how consistent is the response between organ vs. whole-tree level.
2. We examined phenotypic variability for 16 functional leaf (dimensions, nutrient, chlorophyll) and wood traits (density) across two soil types, Ferralitic Soil (FS) vs. White Sands (WS), on two sites for 70 adult trees of *Cecropia obtusa* Trécul (Urticaceae) in French Guiana. *Cecropia* is a widespread pioneer Neotropical genus that generally dominates early successional forest stages. To understand how soil types impact resource-use through the processes of growth and branching, we examined the architectural development with a retrospective analysis of growth trajectories. We expect soil types to affect both, functional traits in relation to resource acquisition strategy as already described at the interspecific level, and growth strategies due to resource limitations with reduced growth on poor soils.
3. Functional traits were not involved in the soil response, as only two traits-leaf residual water content and K content-showed significant differences across soil types. Soil effects were stronger on growth trajectories, with WS trees having the slowest growth trajectories and less numerous branches across their lifespan.
4. The analysis of growth trajectories based on architectural analysis improved our ability to characterise the response of trees with soil types. The intraspecific variability is higher for growth trajectories than functional traits for *C. obtusa*, revealing the complementarity of the architectural approach with the functional approach to gain insights on the way trees manage their resources over their lifetime. Soil-related responses of *Cecropia* functional traits are not the same as those at the interspecific level, suggesting that the effects of the acting ecological processes are different between the two levels. Apart from soil differences, much variation was found across sites, which calls for further investigation of the factors shaping growth trajectories in tropical forests.

## Introduction

Trait-based community ecology seeks to predict the processes of assemblage and maintenance of plant communities over time and space (McGill et al. 2006). The key questions in this field are (i) the identification of ecological processes determining community composition (McGill et al. 2006; Shipley et al. 2016), and (ii) the role of intraspecific variability (ITV) in community assemblages, and to what extent ITV can be ignored by using species-level functional trait means (Violle et al. 2012; Shipley et al. 2016). Trait-based approaches have improved our understanding of the role of ecological processes in community assemblage. Environmental filtering drives community assemblage through the interaction of individuals with the abiotic environment (Kraft et al. 2015): Physiologically challenged individuals are eliminated, so that the breadth of functional trait values is predicted to be small (i.e. functional trait under-dispersion) within local communities. Another process, niche differentiation is based on the interaction of neighbouring individuals, and incorporates the effects of both resource competition and shared predators (Uriarte et al. 2004). For species co-existence, they cannot share exactly the same niche, such that evenness of functional trait value distribution is predicted to be high, leading to functional trait over-dispersion within local communities. Both ecological processes, environmental filtering and niche differentiation, have been demonstrated for various habitats and landscapes, with environmental filtering tending to be more pervasive (Kraft et al. 2008; Swenson and Enquist 2009; Paine et al. 2011; HilleRisLambers et al. 2012).

ITV has long been ignored, or at least underestimated, in trait-based community ecology (Violle et al. 2012; Shipley et al. 2016). This has been the case for studies investigating ecological processes of community assemblages (Schamp et al. 2008; Kraft et al. 2008; Swenson and Enquist 2009); but see Paine et al. (2011). ITV may allow a species to thrive in several communities. First, displaying a large ITV would allow a species to fit a large abiotic spectrum, since there is a higher probability that the required functional trait values compatible with the habitat fall into the possible range of functional trait values of the species. In such a case, the success of individuals in terms of environmental filtering is promoted. Second, displaying a large ITV allows different ways to avoid functional trait similarity with neighbours, and contributes to the niche differentiation among habitats. Recent efforts for incorporating ITV into trait-based community ecology have validated these hypotheses (Fridley and Grime 2010; Jung et al. 2010, 2014; Lepš et al. 2011; Kraft et al. 2014). Moreover, studies have demonstrated that incorporating ITV increases the predictive power of models for species interactions, trait-environment relationships, and ecosystem productivity (Jung et al. 2010; Paine et al. 2011).

Generalist species are particularly useful to understand the role of ITV and environmental filtering in the structuring of community composition. Generalist species are defined here as species able to thrive within a larger range of abiotic conditions than most of species, and generally regarding one kind of condition, i.e. topographic, edaphic, light… First, generalist species tend to display large ITV, and by definition they inhabit large ecological spectrums (Sides et al. 2014). Generalist species offer the opportunity to test hypotheses regarding how ecological processes act at the intraspecific level, how functional traits are mediated regarding the ecological processes, and if they do so in the same way than at the interspecific level. Second, better knowledge on how ITV of functional traits is structured should help to better decide if and how ITV must be accounted for in trait-based community ecology, especially for generalist species, which are often regionally widespread and abundant (Holt et al. 2002; Borregaard and Rahbek 2010; Boulangeat et al. 2012).

The Amazon rainforest has been a rich study field for investigating key questions on trait-based ecology, such as relationships of functional traits with environmental gradients (Kraft et al. 2008). Edaphic gradients have been particularly studied to disentangle drivers of spatial distribution of species and functional traits over the Amazon basin (Sabatier et al. 1997; Clark et al. 1999; Stropp et al. 2011; Allié et al. 2015). The contrast between white-sand (WS) versus ferralitic soils (FS) has been repeatedly used for explaining Amazonian spatial species diversity turnover. WS are mainly quartz soils, representing 3% to 5% of soils in the Amazon basin and exist as island-like spots in a matrix of other soils such as FS (Adeney et al. 2016; Fine and Baraloto 2016).WS are poor in mineral nutrients and acidic, with low-usable water reserves and poor nitrogen mineralization, especially in comparison with common FS. The environmental filtering ensued by the FS-WS heterogeneity has strong impacts on species distribution patterns at the community level (Stropp et al. 2011; ter Steege et al. 2013; Daly et al. 2016; Fine and Baraloto 2016), as well as affecting functional traits. WS flora is characterised by a convergence towards a conservative functional strategy of nutrient acquisition because of the scarce nutrient availability and severe water stress (Grubb and Coomes 1997; Patiño et al. 2009; Fyllas et al. 2009; Fine et al. 2010; Fortunel et al. 2012; Fortunel, Paine, et al. 2014; Fortunel, Ruelle, et al. 2014; Fine and Baraloto 2016). These imply higher leaf mass area (LMA), higher wood density, smaller seeds, and lower leaf nutrient contents associated with higher nutrient use efficiency (Fine and Baraloto 2016), in comparison with other soil types such as FS.

Many generalist species are able to grow on either end of the FS-WS gradient (Fine and Baraloto 2016). A pervasive soil response has been demonstrated for the tree species *Protium subserratum* Engl. (Burseraceae), where Fine et al., found significant differences on the chemical traits associated to herbivory resistance between individual growing on FS and WS, paralleling the environmental filtering acting at the interspecific level (Fine et al. 2013). However, we do not know how more commonly used, morphological, and easy-to-measure functional traits (e.g. LMA, leaf area, leaf thickness, wood density…), largely used in trait-based plant community ecology, vary at the intraspecific level between FS and WS, and if the environmental response of these functional traits mirrors the environmental filtering acting at the community level (Fortunel, Paine, et al. 2014).

Phenotypic adjustment to abiotic factors could occur in different manners according to the plant compartment (i.e. roots, trunk, or leaves; Paine et al. 2011, Freschet et al. 2018, Fortunel et al. 2014) or the function (assimilation, mechanical stability, conduction…; Freschet et al. 2018). For instance, functional traits associated to resource acquisition (e.g. leaf and root traits) could vary independently of functional traits related to resource use (e.g. growth, defense; Fine et al. 2006, 2013). Easy-to-measure, organ-level, functional traits commonly used in trait-based ecology are appropriate to capture a snapshot image of the resource-acquisition strategy (Baraloto et al. 2010). They are generally assumed to be proxies of the individual performance, and therefore to indirectly impact fitness (Violle et al. 2007). But functional traits generally measured in trait-based ecology fail to take into account the growth strategy, which integrates the long-term response of the individual to its environment. Limiting habitats, such as WS, are physical boundaries in terms of available energy, water and nutrients per unit of time for a given plant. Even if functional traits associated to resource acquisition do not vary, the resource scarcity could have an effect on how the tree develops in space and time throughout the whole tree lifespan. Here, we combine functional trait approaches with a whole-tree developmental approach based on retrospective analysis to gain complementary aspects of tree phenotypic responses.

With a whole-tree developmental approach, we can consider the development of the trunk for instance, described as a sequence of repetitive elementary units (e.g. internode, growth unit, annual shoot), universal for vascular plants, and the accumulation and fluctuation of growth, branching, and flowering processes through a tree’s lifespan (Heuret et al. 2006; Guédon et al. 2007; Taugourdeau et al. 2012). Therefore, the accumulation of growth and branching over time can be expressed as a growth trajectory, and represents the ability of trees to develop and produce biomass. Such growth trajectory can be seen as a performance trait, as growth is one of the three main components of individual performance directly impacting fitness (Violle et al. 2007). Moreover, the analysis of the fluctuation of elementary units (internode length, annual shoot length, number of branches…) through tree’s lifespan conveys complementary insights on the determinants of variation of growth trajectory across trees, and further help to characterize different growth strategies. Here, we aim to elucidate the role of ITV in functional traits and growth patterns in allowing species to thrive in different environments by studying the Amazon rainforest genus Cecropia, composed of hyperdominant pioneer tree species, critical in the recovery of Amazon forests.

We focus on *Cecropia obtusa* Trécul (Urticaceae), a widespread Guiana shield generalist species, capable of growing on both FS and WS, and displaying perennial growth marks, which allow for an analysis of life history based on architecture analysis (growth, branching, flowering) through time, making *C. obtusa* a model species for tree architecture and growth (Heuret et al. 2002; Zalamea et al. 2008; Mathieu et al. 2012; Letort et al. 2012). We measured commonly used functional leaf and wood traits, coupled with the growth trajectory (i.e. fluctuation and accumulation of growth over time) and architectural development (i.e. integration of growth, branching, and flowering processes) analyses for *C. obtusa* individuals from two sites with both soil types in French Guiana. We aim to answer the following questions:

i. Is the effect of environmental filtering on functional traits the same at the intraspecific and interspecific levels?
ii. Do the measured functional traits and performance traits response equivalently to soil types for *C. obtusa*?

## Methods

### Terms and definitions

In this study, we use the term of “functional trait” according to the definition of Violle et al. (2007), as any morphological, physiological, or phenological trait which impact fitness indirectly via their effects on growth, survival, or reproduction. But in this study, functional traits specifically refer to easy-to-measure traits, generally measured at the organ level, generally measured in trait-based ecology, and sometimes referred as soft traits (Violle et al. 2007): e.g. leaf area, leaf mass area, wood density… These traits are generally measured at a specific given moment of the plant’s life, disconnected from the developmental trajectory, and ignoring potential ontogenetic effects on the trait value. That is why we oppose functional traits to performance traits in our study. Performance traits are defined here as morphological traits directly related to growth, branching, and flowering processes, and that can be expressed as longitudinal data, i.e. trajectory: internode length, annual shoot length, number of branches per annual shoot, number of inflorescence per annual shoot… according to plant height, or plant age, or node ranking. We also used whole-tree-level traits, defined as traits capturing whole-tree features of architecture at a specific given moment of the plant’s life, such as tree height, the total number of branches and inflorescences, the number of branching orders… Such traits are generally harder to measure than soft traits we refer as functional traits in our study, and are not expressible as longitudinal data as our performance traits. The goal of the use of this specific terminology in the context of our study is to contrast the architectural approach and related measurements of architectural-feature trajectories, which are not so common in trait-based ecology.

### Study species: Why C. obtusa is an appropriate tree model species?

*C. obtusa* has several characteristics that allow the retrospective construction of a tree’s past growth. The growth of *C. obtusa* is continuous (no cessation of elongation) and monopodial (no death of meristem), the tree is made of a set of axes, where each one is composed of an ordered, linear, and repetitive succession of phytomers (i.e. the set of a node, an internode, a leaf, and its axillary buds; Fig. S1). Leaves are stipulated, with an enveloping stipule named calyptra which has a protective function (Fig. S1). At the leaf establishment, the calyptra sheds leaving a characteristic ring scar delimiting the associated internode, and usable as a permanent growth marker. The 10-day stable phyllochron (i.e. rhythm of leaf production) associated with such permanent growth marker allows for the retrospective analysis of tree growth and development, covering the tree’s lifespan (Heuret et al. 2002; Zalamea et al. 2012).

There are three lateral buds in the axil of each leaf (Fig. S1). The central bud is vegetative and can develop into a new axis. The two others are proximal lateral buds of the vegetative central one and can develop into inflorescences The inflorescences leave permanent scars after shedding, allowing the retrospective analysis of tree’s lifespan flowering events. The same retrospective analysis is possible with branching events since the presence of past branches remains visible.

### Study site

Two sampling sites were selected in French Guiana: (1) Counami, along the Counami forestry road (N5.41430°, W53.17547°, geodesic system WGS84); and (2) Sparouine, along the national road 5 (N5.27566°, W54.20048°). The warm and wet tropical climate of French Guiana is seasonal due to the north-south movement of the Inter-Tropical Convergence Zone. Annual rainfall is 3,041 mm year-1 and annual mean air temperature is 25.7 °C at Paracou experimental station (Gourlet-Fleury et al. 2004) situated nearly at 30 km and 150 km to the east of Counami and Sparouine sites respectively. There is one long dry season lasting from mid-August to mid-November, during which rainfall is < 100 mm month-1. The two studied sites (Counami and Sparouine) are characterised by rainfall differences (Fig. S2). Counami shows higher levels of rainfall and higher contrasts between the long rainy and the long dry seasons. For each of the two sites, two micro-localities are identified corresponding to two soil types: ferralitic soils (FS) and white-sand soils (WS). Local sites were chosen to be well drained and on upper slopes.

### Plant material, study conception, and sampling

Individuals had grown in clearings and formed a secondary forest where they are the dominant species together with *C. sciadophylla*. A total of 70 trees were selected in September and December 2014 respectively for Counami and Sparouine sites: 32 in Counami and 38 in Sparouine. Soil samples were taken at the same time for pedological analysis. On the Counami site, where individuals are widely spaced, a soil sample was taken at the basis of each individual tree. On the Sparouine site, where individuals where clustered, 9 soil samples were taken, as each soil sample was representative of 4-6 individuals located no further than 30m from the soil sample spot.

As *C. obtusa* is dioecious, only pistillate (i.e. female) trees were felled to avoid potential sex-related variability in the measured functional traits. Trees were not felled according to the same scheme in the two sites. Trees were preselected to have as close as possible comparable diameters at breast height (DBH), and age was estimated with binoculars according to the method described by Zalamea et al. (2012). By counting the number of internodes we were able to estimate the age of trees as each internode is produced in 10 days (Heuret et al. 2002; Zalamea et al. 2012). In Sparouine, all individuals correspond to a single colonisation pulse on both soil types: all individuals have similar age (7-10 years), with DBH of 11.94 to 25.70 cm, and heights of 13.85 to 23.20 m (Fig. S3). Both soil types were represented by 19 individuals and all individuals were felled and measured between the 14th and the 19th of September 2015. Thus, season-, size-, and age-related effects on functional traits are controlled for soil and individual comparisons.

The experimental design at Counami was different. The forestry road was opened gradually, and therefore the age of the trees differed according to the road section (Zalamea et al. 2012). All individuals assigned to WS at Counami were selected at a single small WS patch located 6 km after the entrance of the road. WS trees represented a single colonisation pulse and were of similar age (14-16 years, except one significantly older with 22.8 years old), with DBH from 6.21 to 15.18 cm, and heights from 10.27 to 16.18 m, (Fig. S3). It was not possible to choose trees on FS on a single restricted area because of the perturbation of soil structure by the logging machines and because we excluded trees on down slopes. Consequently, FS trees were sampled between km 6 to 11 of the forestry road and included different cohorts with different ages (7-23 years), DBH of 9.55 to 22.44 cm, and heights of 12.16 to 22.63 m (Fig. S3). Thirteen and nineteen individuals were sampled on FS and WS respectively. Counami trees were felled at different dates, from September 2014 to April 2016. The contrasted protocol was chosen to study seasonal and ontogenetic effect on leaf traits, but the results of such analysis will not be addressed here. No seasonal effects on leaf traits were detected, and ontogenetic effects on functional trait were standardised, as presented in the Statistical analyses part.

### Soil properties

Pedological analyses included granulometry, moisture content, pH, organic matter content, and contents of exchangeable cations (Appendix S1, with detailed abbreviations). The complete sampling procedure is described in the Appendix S1. Exchangeable cations were analysed divided by cation-exchange capacity (CEC) to avoid correlations between the former and the latter. We also calculated a soil index of fertility as: soil_index_ = (K+Ca+Mg+Na)/CEC.

The a priori classification of soil types (FS-versus-WS) was confirmed by pedological analyses of the soil properties within each site. The described pattern of soil properties is congruent with that reported in the literature (Adeney et al. 2016; Fine and Baraloto 2016). WS consist of a large proportion of coarse sand with high Ca:CEC (calcium on CEC) and C:N (carbon on nitrogen) ratios. FS consist of a large proportion of clay and silt with high moisture, N, C, MO P_tot_ (total potassium) contents and a high Al:CEC (aluminium on CEC) ratio. Based on water availability, N content, and soil_index_, the site fertility can be ordered as COU-FS > SPA-FS > COU-WS = SPA-WS. Sparouine WS are characterised by higher H:CEC and Fe:CEC ratio than Counami WS. The related results are presented in Appendix S1. Within sites, the difference between soil types is more contrasted in Counami than in Sparouine.

### Architectural and functional traits

For all individuals, we measured a suite of performance and whole-tree-level traits at phytomer and whole-tree levels to characterise growth, branching and flowering dynamics, and the resulting tree architecture. Retrospective analysis of development allows us to consider tree developmental trajectories as growth performance traits (i.e. the height-age relationship). Such approach considers the development of the trunk only (i.e. it does not include the complexity of branching events) described as a sequence of phytomers. Three variables were measured for each phytomer: (1) internode length (2) vegetative bud state coded as: 0 for not developed or aborted; 1 for developed, present or pruned, (3) inflorescence bud state coded as: 0 for no inflorescence; 1 for pruned or present inflorescences. Features for bud states are treated as binary values: presence or absence. As suggested by Davis (1970), Heuret et al. (2002), and Zalamea et al. (2008), we analysed periodical fluctuations in internode length, which are driven by seasonal variations of rainfall (Zalamea et al., 2013), as well as the rhythmic disposition of inflorescences and branches to infer the past development of the tree, and model its growth dynamic (section statistical analysis and Appendix S2).

As a first step, the fluctuation of internode length allowed us to estimate (i) the growth representing a single year as the shortest internodes are associated with the peak of the dry season, (ii) the age in days after germination of any internode along the trunk, and (iii) the yearly average time taken by the tree to produce an internode (i.e. the phyllochron).

As a second step, to understand how the trees undergo changes in growth strategies in the two types of soils, we analysed (i) variations of phyllochron, internode length, and annual shoot length over time, and (ii) contribution of the number of internode vs internode length in the annual shoot length variation (See Appendix S2 for the followed methodology).

As a third step, we analysed how these different potential growth strategies (i.e. number vs. length of internodes) determine the cumulative tree height over time, namely the growth performance. Finally, to study space-foraging performance and reproductive performance we analysed the cumulative branching and flowering over time. The measured and estimated traits presented as longitudinal sequences, are shown in Table 1. Whole-tree-level traits were also measured (Table 1). Functional traits were measured at the leaf level (Table 2) as proxies of leaf resource capture, while trunk wood specific gravity was measured as indicator of stem transport, storage capacity, and mechanical strength (Baraloto et al. 2010). We measured leaf-level traits for only one leaf per individual: either the third or the fourth leaf under the apex of the A1 axis. In this way, potential effects of plant spatial structure and leaf senescence and on variation of leaf-level traits are controlled. Leaf lifespan along the A1 axis was estimated for each tree by counting the number of leaves on a given axis and multiplying it by the known mean phyllochron (10 days, Heuret et al. 2002). The complete sampling procedure for functional traits is described in Appendix S3.

**Table 1.** List of measured growth and tree-level traits.

| Trait | Abbreviation | Unit |
| --- | --- | --- |
| *Performance traits* |  |  |
| Internode length | IN | mm |
| Internode length residuals |  | - |
| Phyllochron |  | day |
| Annual shoot length | AS | cm |
| Number of nodes per AS |  | - |
| Cumulated tree height |  | m |
| *Tree-level traits* |  |  |
| Age after determination | Age | year |
| Tree height | Height | m |
| Diameter at breast height | DBH | cm |
| Number of trunk internodes | IN<sub>A1</sub> | - |
| Number of A2 bearing branches | Br<sub>bear</sub> | - |
| Number of A2 dead branches | Br<sub>dead</sub> | - |
| Total number of A2 branches | Br<sub>tot</sub> | - |
| Branching order | Order | - |
| Total number of inflorescences | Fl<sub>tot</sub> | - |
| Total number of leaves | Leaf<sub>tot</sub> | - |
| Total estimated crown area | A<sub>crown</sub> | m<sup>2</sup> |
| First branching height | Br1stH | m |
| First flowering height | Fl1stH | m |
| First branching node rank | Br1stIN | - |
| First flowering node rank | Fl1stIN | - |
| First branching age | Br1stAge | year |
| First flowering age | Fl1stAge | year |

**Table 2.**
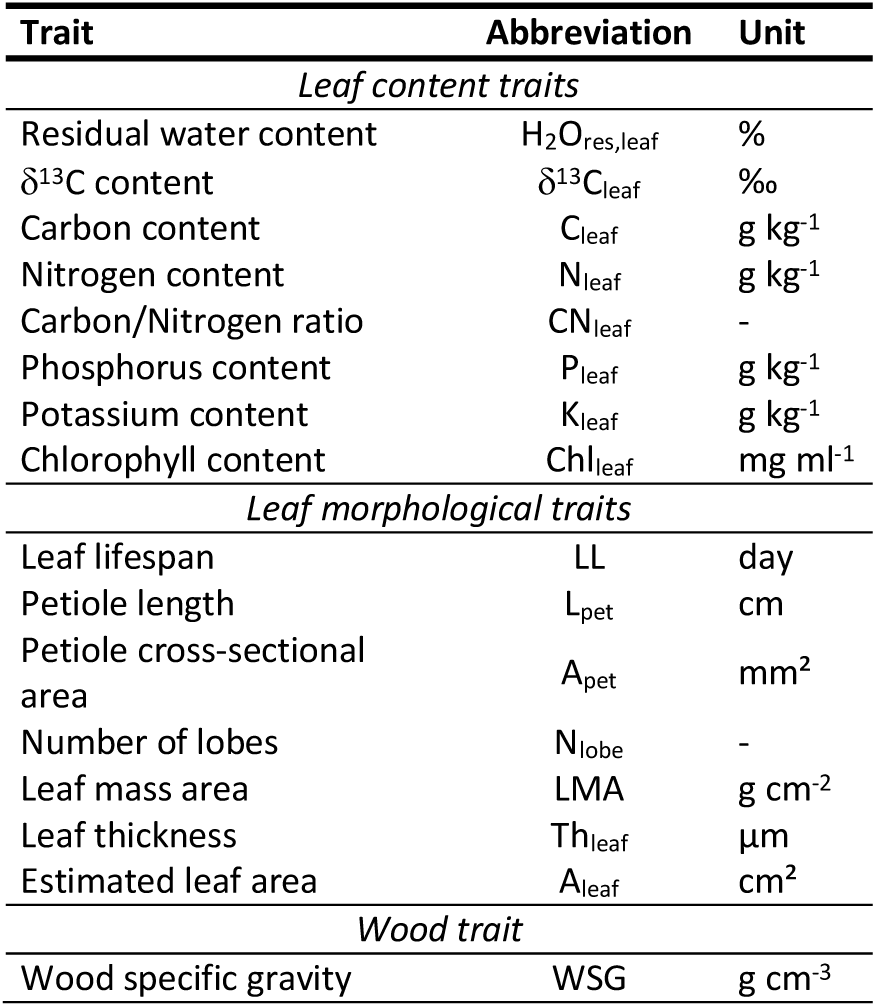
List of measured functional traits.

### Statistical analyses

Topology of trees and the different pedological, whole-tree-level, and functional features associated with each repetitive unit are coded in sequences in Multi-scale Tree Graph format (MTG; Godin & Caraglio, 1998; Godin, Costes, & Caraglio, 1997).

Statistical analyses relative to developmental trajectories were conducted with AMAPmod (op. cit), now integrated in the OpenAlea platform, re-engineered and named ‘VPlants’ (Pradal et al. 2013), and the R programming language (R Core Team 2018). We conducted autocorrelation coefficients on internodes (length, branch presence, inflorescence presence) to confirm an annual periodicity at stand level (i.e. soil x site) for growth, branching, and flowering processes. Methods and results regarding the analysis of autocorrelation coefficients are presented in Appendix S4. To analyse fluctuations of internode length, we used a method of time series analysis relying on a decomposition principle of signals, described as follows. The different sources of variation, such as long-term changes at low-level frequency (i.e. over hundreds of internodes and several years) vs short-term changes at high-level frequency fluctuations (i.e. over tens of internodes and few months), are identified and filtered (Guédon et al. 2007). Firstly, we calculated a moving average to extract the trend of internode length sequences in a similar way as Zalamea et al. (2008). Having extracted the trend, we looked at local fluctuations by examining the residuals. Residuals were generated by dividing for each internode, its length by its moving average (Appendix S2 for details). The analysis of residuals allowed the identification of the limits of the long dry season in September/October for successive years, since shorter internodes are elongated during this period as shown for *C. obtusifolia* Bertol. (Davis, 1970), *C. peltata* L., and *C. sciadophylla* Mart (Zalamea et al. 2013). Delimitation of annual growth for each individual allowed the estimation of a mean phyllochron for each year according to the node rank (Appendix S2). Knowing the phyllochron allowed the conversion of the rank node to a temporal scale, namely the age. Finally, by considering the length or the number of nodes elongated between two successive dry seasons, we estimated the annual shoot length (Table 1). Growth strategies are studied as (i) variations of phyllochron, internode length, and annual shoot length over time, and (ii) contribution of the number of internode vs. internode length in the annual shoot length variation. Significant differences in performance traits (i.e. internode length, phyllochron, AS length, number of internodes per annual shoot) between FS and WS were identified based on a confidence interval at 95% around the mean trajectory of the considered performance trait. A mean trajectory was calculated and plotted for each soil type within each site.

To test the effect of soil type on the variability of growth trajectories –which are longitudinal data by nature-, we tested the correspondence of distribution of (i) soil types, with that of (ii) clusters defined by statistical signatures of growth trajectories. The clusters were characterised with a clustering method on the generated longitudinal data (Table 1), with the kml R package (Genolini and Falissard 2009). It is a classification method based on an implementation of “k-means”, itself based on a minimization function of distances among trajectories. For each trait, 100 simulations were used, and decisions are based on the Calinski-Harabasz criterion. The optimal number of clusters corresponds to a maximisation of the Calinski-Harabasz criterion. The dependency of defined clusters on soil types is evaluated with a Pearson’s chi-squared test. Analyses relative to soil, whole-tree-level, and functional trait data are realised in R language.

Unlike the trees sampled at Sparouine, Counami trees formed a non-even-aged population that we sampled at different moments of the year. We tested by multiple regression analysis for the potential effect of season as well as the effect of ontogenetic stage of individuals, assessed by the age of the tree, on the leaf traits of all 70 sampled leaves. Overall, no seasonal effect on leaf functional traits was found. Ontogenetic effects were found for some functional traits and were taken into account before testing for soil effects. Principal Component Analysis (PCA) on soil properties and functional traits were conducted with the ade4 (Chessel et al. 2004) and Factoextra (Kassambara and Mundt 2016) R packages. For the PCA analysis, when ontogenetic effects were found on a given functional traits, residuals of the linear model between this trait and tree age were used. The effect of soil on functional and whole-tree-level traits was tested with linear mixed-effect models (LMER), with the soil gradient modelled by tree coordinates along the first axis (45,4%) of the soil PCA (Appendix S1). Soil and tree age –if a tree age effect was detected for a given functional trait-were set as fixed effects, and site as a random effect. A comparison of factorial coordinates of individuals was conducted for each axis based on a nested-ANOVA and a post-hoc Tukey’s HSD test.

## Results

### Developmental approach: architecture and growth trajectory

Fig. 1 shows significant differences in trajectories of performance traits between FS and WS, based on plotted confidence intervals around the mean trajectory. Internode length was significantly shorter for WS in comparison to FS in Counami (Fig. 1c) for the first 5 years only. These first 5 years corresponded to the ontogenetic stage with the longest internodes. No difference in internode length was found in Sparouine between FS and WS (Fig. 1e). Clusters of internode length trajectories significantly matched soil type distributions in Counami (P < 0.01), but not in Sparouine (P > 0.05; Fig. 1d,f). Phyllochron –and the related variable, the number of nodes per annual shoot-, were not significantly different between FS and WS for either site (Fig. 1a; Fig. S4a,c). Clusters of the trajectories of the number of internodes per annual shoot trajectories significantly matched soil type distributions in Sparouine (P < 0.01), but not in Counami (P > 0.05; Fig. S4b,d), based on Pearson’s chi-squared tests. Annual shoot length was significantly shorter for WS in comparison to FS in Counami (Fig. S4e) for the first 5 years only. No difference in annual shoot length was found in Sparouine between FS and WS (Fig. S4g). Clusters of annual shoot length trajectories significantly matched soil type distributions in Counami (P < 0.01), but not in Sparouine (P > 0.05; Fig. S4f,h), based on Pearson’s chi-squared tests.

**Figure 1.**
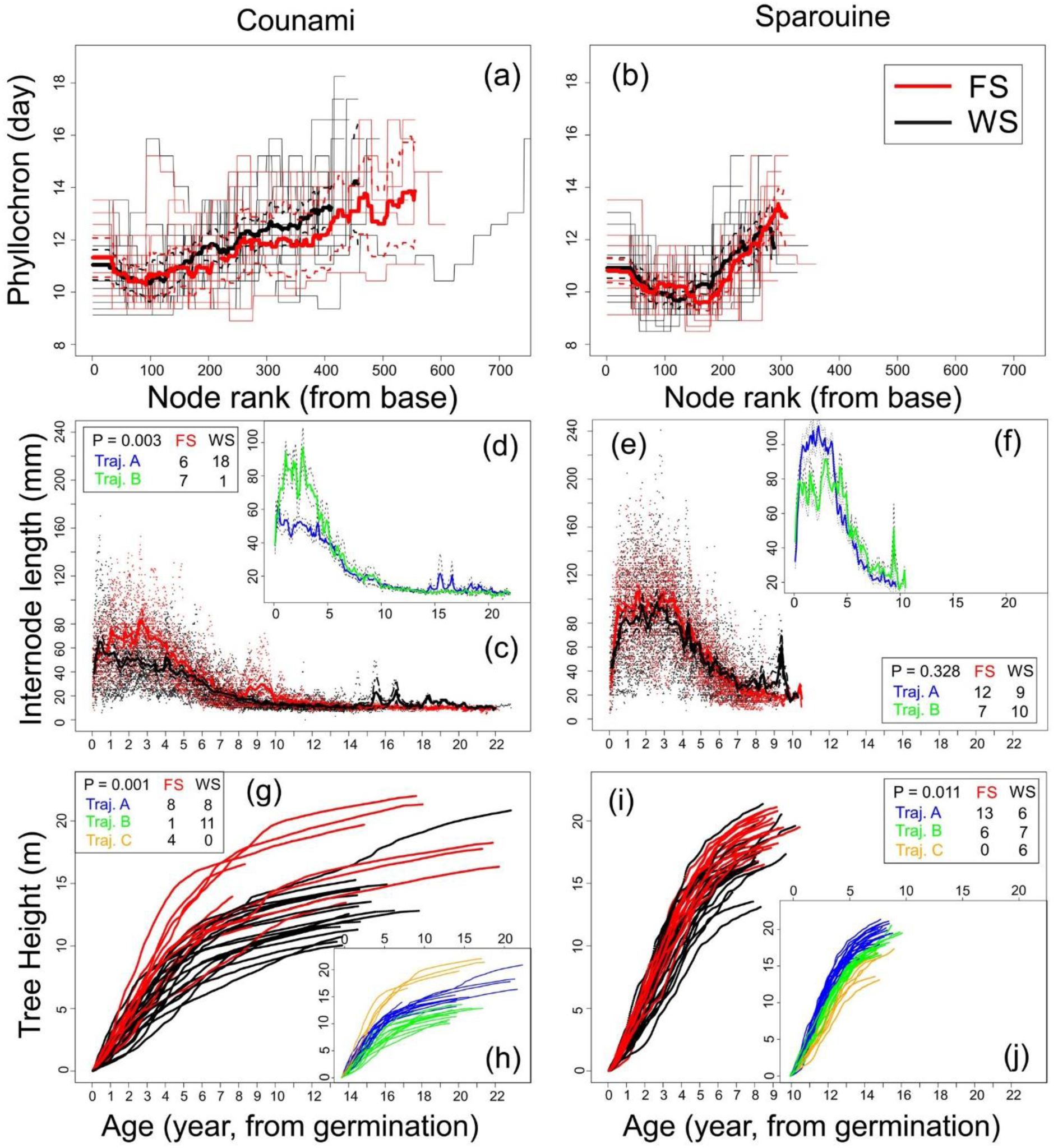
Architectural growth traits according to node rank and age (years). Main boxes represent features according to soil types. Inboxes represent mean trajectories after clustering longitudinal analyses (*kml*). The left column represents Counami trees, the right column represents Sparouine trees. Distributions between soil types and kml-trajectories are represented with Pearson chisquared test. Red: ferralitic soils; black: white-sand soils. Blue: trajectory A; green: trajectory B; orange: trajectory C. Thick lines: means; dashed lines: confidence intervals at 95%.

For both sites, there was a pattern for FS trees to be higher than WS trees for a given age (Fig. 1g,i). For both sites, it was possible to identify two main growing phases. The phases were differentiated by variations in growth rates over the tree’s lifespan. The first phase covered the first 5-7 years, except for FS Counami trees where it was the first 9-10 years. The second growing phase was defined by a slower growth rate, which remained constant for all individuals. For both sites, cluster of tree height trajectories significantly matched soil type distribution based on a Pearson’s chi-squared test (P < 0.05; Fig. 1h,j).

The analysis of the cumulated number of pairs of inflorescences on the trunk indicated that there was no significant difference between FS and WS for both sites based on confidence intervals (Fig. 2a, b). In Counami trees, there was a significant difference in the cumulated number of branches of the trunk between FS and WS after 5-6 years old (Fig. 2c). In Sparouine trees there was no significant difference in the cumulated number of branches on the trunk between FS and WS (Fig. 2d).

**Figure 2.**
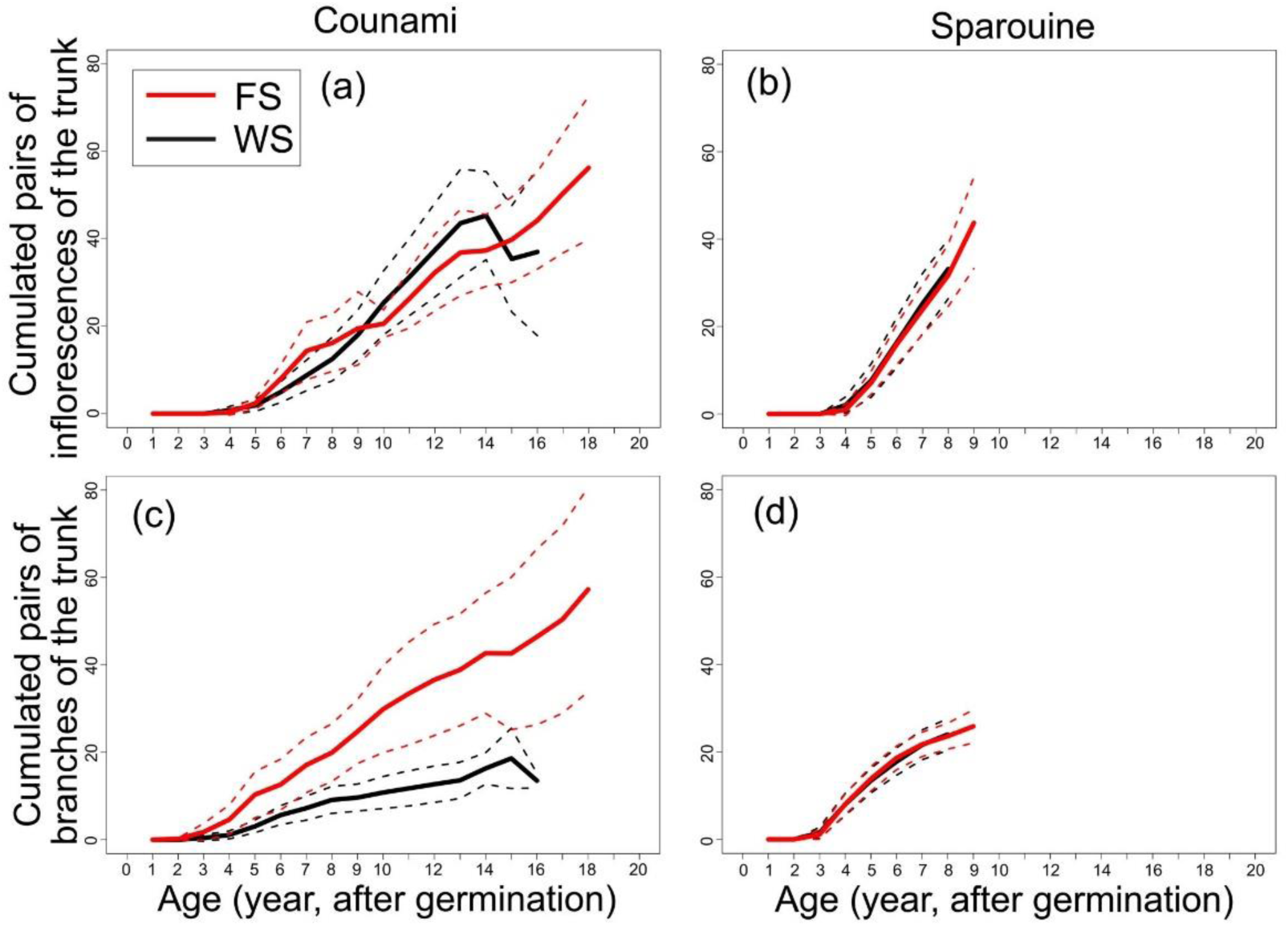
Cumulated number of trunk nodes with pairs of inflorescences and number of branches per annual shot according to the age (year). (a) and (b) Means for inflorescences for Counami and Sparouine respectively. (c) and (d) Means for branches for Counami and Sparouine respectively. Red: ferralitic soils; black: white-sand soils. Thick lines: means, dashed lines: confidence intervals at 95%.

A significant effect of soil was identified for tree height, DBH, the branching order, and the height of the first flowering and first branching (P < 0.05; Table 3; LMER), with all whole-tree-level traits increasing in FS.

**Table 3.** Linear mixed-effect models for measured architectural dimension traits between soil types and sites.

| Abbreviation | Unit | Age | Soil | Direction | Counami FS | Counami WS | Sparouine FS | Sparouine WS | Range |
| --- | --- | --- | --- | --- | --- | --- | --- | --- | --- |
| Height | m | *** | *** | + | 16.13 ± 1.92 | 13.31 ± 1.10 | 18.98 ± 0.65 | 17.27 ± 0.99 | 10.05 – 21.99 |
| DBH | cm | *** | ** | + | 15.02 ± 2.24 | 12.44 ± 2.33 | 19.30 ± 0.97 | 18.24 ± 1.52 | 6.21 – 30.49 |
| IN<sub>A1</sub> | - | *** |  |  | 429.2 ± 87.0 | 457.9 ± 35.0 | 289.5 ± 10.1 | 277.9 ± 14.8 | 221 – 765 |
| Br<sub>bear</sub> | - |  | * | + | 7.85 ± 1.75 | 4.32 ± 1.55 | 10.74 ± 1.79 | 11.11 ± 1.82 | 0 – 19 |
| Br<sub>dead</sub> | - | *** |  |  | 10.23 ± 6.02 | 7.47 ± 1.97 | 12.84 ± 2.35 | 10.58 ± 2.81 | 0 – 33 |
| Br<sub>tot</sub> | - |  |  |  | 13.62 ± 4.07 | 6.79 ± 2.90 | 21.47 ± 3.80 | 24.16 ± 8.17 | 1 – 75 |
| Order | - | *** | * | + | 2.92 ± 0.41 | 2.32 ± 0.26 | 3.16 ± 0.17 | 3.26 ± 0.20 | 1 – 4 |
| Fl<sub>tot</sub> | - |  |  |  | 55.54 ± 17.11 | 46.42 ± 12.31 | 75.68 ± 24.95 | 121.9 ± 68.0 | 0 – 657 |
| Leaf<sub>tot</sub> | - |  |  |  | 140.6 ± 37.6 | 75.74 ± 31.54 | 170.74 ± 31.35 | 200.3 ± 72.7 | 25 – 674 |
| A<sub>crown</sub> | m<sup>2</sup> |  |  |  | 16.44 ± 6.11 | 6.893 ± 3.472 | 21.79 ± 4.17 | 26.78 ± 7.89 | 1.656 – 74.10 |
| Br1stH | m |  | ** | + | 9.04 ± 0.80 | 8.20 ± 0.80 | 10.45 ± 0.50 | 9.05 ± 0.85 | 5.27 – 14.56 |
| Fl1stH | m | * | *** | + | 11.29 ± 1.38 | 9.12 ± 0.89 | 18.67 ± 0.71 | 16.71 ± 0.94 | 6.62 – 20.99 |
| Br1stIN | - |  |  |  | 141.8 ± 22.6 | 178.7 ± 32.1 | 117.0 ± 6.8 | 113.7 ± 8.6 | 81 – 358 |
| Fl1stIN | - | * |  |  | 188.6 ± 26.3 | 198.6 ± 32.7 | 284.2 ± 7.9 | 264.1 ± 13.9 | 101 – 412 |
| Br1stAge | year |  |  |  | 4.272 ± 0.752 | 5.297 ± 1.053 | 3.358 ± 0.220 | 3.263 ± 0.275 | 2.332 – 11.94 |
| Fl1stAge | year |  |  |  | 5.709 ± 0.821 | 8.833 ± 0.986 | 8.307 ± 0.231 | 7.699 ± 0.406 | 3.132 – 11.49 |

### Characterisation of functional traits

The first (28.5 %) and second axes (18.5 %) of the PCA for functional traits explained 47.0 % of the inertia (Fig. 3a). The first axis (28.5 %) was driven by C:N_leaf_, L_pet_, A_pet_, A_leaf_ and N_leaf_. The second axis (18.5 %) is driven by H2O_res,leaf_, and K_leaf_. Conditions (i.e. soil types and sites) were not differentiated along the first axis (Fig. 3b; P > 0.05; ANOVA), but differentiated along the second axis (P < 0.001; ANOVA) with Counami trees differing from Sparouine trees. Significant effect of soil was detected for leaf residual water content and leaf K content (P < 0.05; Table 4), with lower residual water content but higher K content for FS trees.

**Table 4.**
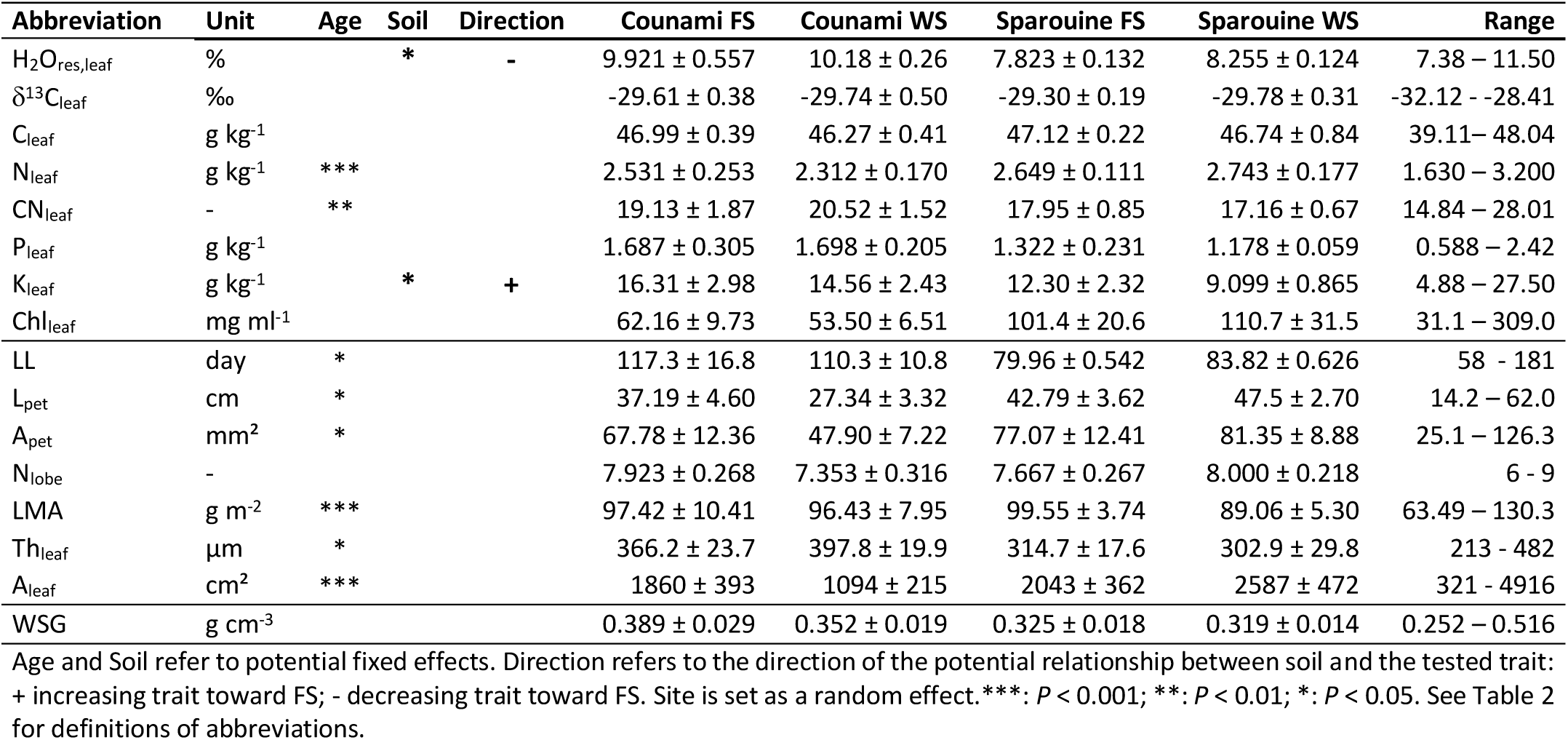
Linear-mixed effect models for measured functional traits between soil types and sites.

**Figure 3.**
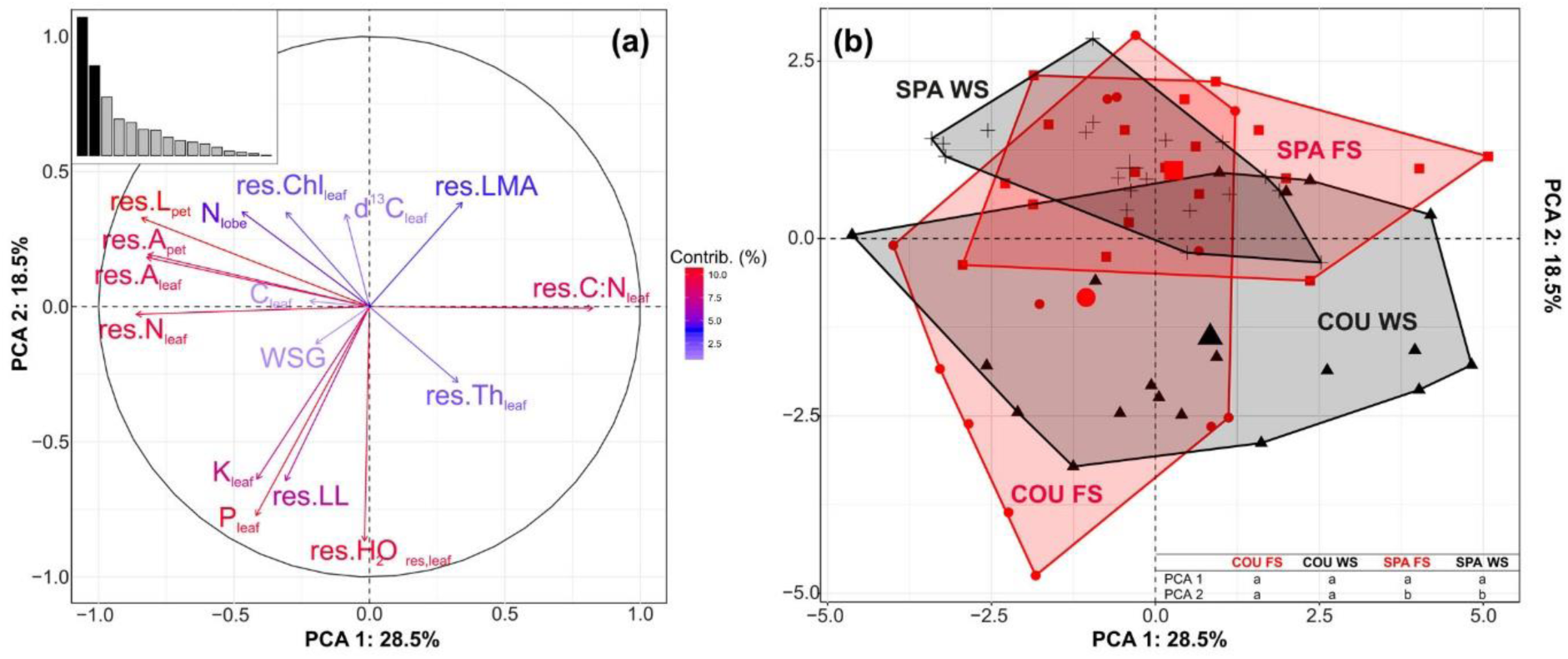
Principal component analysis (PCA) on functional traits for the two sites for 70 trees. (a) Correlation circle of data with the histogram of inertia. (b) Individual factor map of data according to soil types and sites. In (a), the colour gradient indicates the contribution of each variable to the axis. See Table 3 for definitions of abbreviations. “Res” prefixes indicate residuals after removing the ontogenetical effect. In (b), significant differences in coordinates (P < 0.05; ANOVA) between soil types and sites are indicated by letter according to the considered axis. Red: ferralitic soils; black: white-sand soils; COU: Counami; SPA: Sparouine.

## Discussion

To our knowledge, our study is the first incorporating both, tree architectural development and functional traits, in relation with the environment. It is also the first to quantify functional traits for an Amazonian generalist species regarding soil types: FS and WS. The pedological analysis confirmed strong contrasts in soil characteristics between FS and WS, opening the possibility of soil-related phenotypic response. Only two functional traits were responsive to soil type. But they were not the traits known to be the most structuring in the main economic spectra identified (e.g. LMA for the leaf economic spectrum –Wright et al., 2004- and wood density for the wood-economic spectrum –Chave et al., 2009; Zanne et al., 2010-). Here, our integrated approach, combining functional trait and architectural development, showcases how environmental constraints can impact differently on 1) the response of traits between the organ level and the resource-acquisition axis, and 2) the whole-tree level and the resource-use axis, at least at the intraspecific level. Our study also demonstrated that environmental constraints can also have different effects on traits at the intraspecific and the interspecific level.

### Soil-response of functional traits is not the same between intra- and interspecific levels

At the community level in the Amazon rainforest, edaphic contrasts lead to strong environmental filtering mediated by functional traits. WS flora is characterised by a convergence in functional traits, particularly towards a conservative strategy (Grubb and Coomes 1997; Patiño et al. 2009; Fyllas et al. 2009; Fine et al. 2010; Fortunel et al. 2012; Fortunel, Paine, et al. 2014; Fortunel, Ruelle, et al. 2014; Fine and Baraloto 2016). It implies high LMA, high WSG, and low leaf nutrient contents associated with high water use efficiency (i.e. high photosynthetic assimilation to stomatal conductance ratio) for WS tree species (Fine and Baraloto 2016), contrary to FS tree species. Fine & Baraloto (2016) show how WS are limiting for plant development, due to resource scarcity, and how resource scarcity in WS is an abiotic factor selecting for functional trait optima diverging from the functional trait optima found on FS. However, the intraspecific *Cecropia* functional trait response to the same environmental heterogeneity did not parallel the environmental filtering operating on functional traits at the community level (i.e. high LMA and WSG on WS, low LMA and WSG on FS…). This indicates that ecological processes, such as environmental filtering and biotic interactions, work in different ways at the interspecific level and the *C. obtusa* intraspecific level.

A striking result is that the soil was not a driver of the variation of measured leaf and wood traits. Only two leaf traits were responsive to soil types: leaf residual water content and leaf K content. The residual water content, which is not a commonly used as functional trait, is indicative of the capacity of leaf tissues to retain water through osmotic adjustments (Bartlett et al. 2012). The residual moisture content was positively correlated to K content (results not shown; P < 0.001; R^2^ = 0.210), which plays a central role in the maintenance of osmotic integrity of cells and tissues (Marschner 1995). Such correlation between residual water content and soil type suggests that edaphic water stress is one of the primary factors underlying the FS-WS gradient, further shaping the phenotypic response, especially for functional traits related to hydraulics and drought tolerance. This is consistent with the pedological analysis which indicates that water availability strongly contributes to the first axis of the PCA performed on soil characteristics, and underlies the FS-WS gradient (Appendix S1).

Such weak functional trait response was unexpected. Three non-mutually exclusive reasons can be explored to explain why only two functional traits responded to changes in soil type.

1. The leaf and wood functional traits we measured can be subjected to strong variation with the succession of tree ontogenetic stages. This ontogenetic effect can be related to changes in local environment with tree growth such as light (Roggy et al. 2005; Coste et al. 2009; Dang-Le et al. 2013). This ontogenetic effect can also be related to mechanical and hydraulic constraints with self-support and long-transport distance (Ryan et al. 2006; Niklas 2007; Oldham et al. 2010; Bettiati et al. 2012; Rungwattana Kanin et al. 2017; Prendin et al. 2018). Two-to-3 fold variation with ontogeny in leaf and wood functional traits has been demonstrated across several studies (Roggy et al. 2005; Coste et al. 2009; Dang-Le et al. 2013; Rungwattana Kanin et al. 2017; Lehnebach et al. 2019). As we consider the trajectory of internode length variation as a performance trait, the trajectory of leaf trait variation with ontogeny can be considered as an integrated functional trait with its own functional significance regarding environmental filtering with soil types. Alternatively, it can be hypothesised that environmental filtering with soil types decisively occurs at seedling and sapling stages, therefore leaf functional traits expressed at these stages would be more responsive.
2. There are relevant functional traits we did not consider in our study. It has been shown that water availability is the leading climate driver of Amazonian rainforest tree growth (Wagner et al. 2012). Water relation and drought-resistance traits, such as drought-induced vulnerability to embolism and stomatal sensitivity, leaf turgor loss point, root depth, crown area to sapwood area ratio, may have played a central role in ensuring growth and survival on the different soil types (Urli et al. 2013; Anderegg et al. 2016; O’Brien et al. 2017; Adams et al. 2017; Eller et al. 2018). Differences in Amazonian soil characteristics can also impact the root system properties (Freschet et al. 2017), including mycorrhizal fungi associations. For instance, it has been shown that ectomycorrhizal mutualisms are much more common on WS (Roy et al. 2016), and several studies suggest that ectomycorrhizal species may be better able to acquire nutrients (Reich 2014).
3. The different plant strategies, or life histories, can be defined along two important strategic axes of plant functioning: the resource acquisition (e.g. photosynthesis, soil nutrient absorption) axis and the resource use (e.g. growth, defense and secondary metabolites) axis (Reich 2014). The functional traits (i.e. leaf and wood traits) measured here are related to resource acquisition, and poorly captured how resources are used. Trees may not necessarily respond to WS resource scarcity by modifying functional traits related to the acquisition axis. But instead, the reduced resources assimilated in a given time may be translated into reduced resource use possibilities. Since plants are organisms with undetermined development, growth remains one of the largest carbon and nutrients sink across lifespan. Thus, growth may be a component of an adaptive response to resource scarcity. Deciphering growth processes and strategies, and quantifying their variations, could represent an opportunity for studying changes along the resource use axis, in relation to the environment.

Regarding biotic interactions, the studies of Fine et al. (2004, 2006) suggest that herbivory pressure could be a primary driver of ecological speciation and diversification within a genus on WS, because of higher costs of tissue lost associated with resource-poor habitats. The resource scarcity selects for structures with long lifespan, and resistant to herbivory pressure. Conversely, *Cecropia* trees are characterised by short lifespan and high growth rates, in relation to their pioneering and competitive strategy, which is in contradiction with a conservative strategy privileging long tree lifespan. Under such hypothesis, selection for light competitiveness would be prevalent on selection for a conservative strategy. This would explain why functional traits such as LMA and WSG are not impacted by soil types as demonstrated by our study. In order to achieve herbivory resistance, three types of defense can be produced: structural, chemical, and mutualistic. Here again, the non-dependence of functional traits such as LMA and WSG on soil types suggests that structural defenses are not required to respond and to allow *Cecropia* trees to grow on WS. Chemical traits related to herbivory resistance have been shown to vary between FS and WS for the generalist tree species *Protium subserratum* (Burseraceae; Fine et al. 2013). Chemical traits related to herbivory pressure, and the herbivory pressure in itself, are poorly known for *Cecropia* trees (Latteman et al. 2014); but functional traits related to herbivory avoidance could play an important role in the strategy required to allow *C. obtusa*’s generalist behaviour (Folgarait and Davidson 1994, 1995; Latteman et al. 2014), and need further investigations. Finally, *Cecropia* species are also famous for their mutualism with the Azteca ant species, where ants offer a protection against visitors by biting (Schupp 1986; Dejean et al. 2009). During field work, we observed ant occupancy on both sites and on both soil types, suggesting an undisturbed interaction between ants and host plants.

### Only performance traits are responsive to soil types, not functional traits

Our results suggest that phenotypic response to soil change is mediated by the architectural development, capturing performance traits related to growth and reproduction, rather than functional traits.

An analysis of growth trajectory based on architectural development analysis is a useful tool for the quantification of the resource use strategy. The autocorrelation function at the stand level confirmed a high degree of periodicity across all individuals for growth, flowering, and branching processes (Appendix S4). With the analysis of internode fluctuations, this periodicity has been shown to be annual, and further allowed to shift on a temporal scale and to conduct our retrospective analysis of architecture. We clearly showed that soil types impacted the overall growth trajectory (i.e. cumulated tree height according to age) for both sites, with WS trees having the lowest trajectories. For any given age, WS trees were always smaller, due to resource scarcity. However, such pattern is less noticeable on Sparouine trees. The site difference could be explained by (i) the less pronounced contrasts between FS and WS in Sparouine as shown by our pedological analysis (Appendix S1), and (ii) the rainier dry season in Sparouine (Fig. S2). Under the assumption that the interaction between WS and water scarcity during the dry season is deleterious for tree growth, this may also explain the generally strongest growth trajectories in Sparouine in comparison to Counami. These substantial site effects on tree phenotype calls for investigating a larger geographic gradient to precise (i) the environmental gradient underlying the geographic gradient (i.e. rainfall, seasonality, irradiance; Wagner et al. 2012), and (ii) the phenotypic response to this environmental gradient.

Regarding the growth strategy, soil type showed a significant effect on both internode length and annual shoot length in Counami, but not in Sparouine. When the soil effect was strong enough, the differences in annual shoot length between soil types corresponded mainly to variations in internode length rather than variations in number of nodes per annual shoot. Reducing the number of nodes per annual shoot would imply the increase of the phyllochron, thus reducing the number of leaves produced per year. Such mechanism would critically affect tree carbon balance, as the contribution of a given leaf to the carbon balance is disproportionate in comparison to most of species: *C. obtusa*’s leaves are large (1,000-5,000 mm^2^, Levionnois et al., data not published) but few (100-600 leaves, Table 3). Similarly, Zalamea et al. (2013) found no difference in phyllochron between *C. sciadophylla* from two distanced locations with contrasting rainfall. The architectural analysis also shows that WS trees in Counami had significantly fewer cumulated branches, and lower branching order, than those in FS. WS trees in Counami have, therefore, reduced space and light foraging capacities, decreasing their competitiveness. Because flowering is synchronous on all crown axes (Heuret et al., 2003), the energetic production cost of inflorescences and seeds is exponentially related to the number of main branches. Therefore, WS trees in Counami must also have comparatively reduced reproductive and dispersive abilities, leading to a reduced overall fitness compared to their FS conspecifics, under the assumption that FS and WS trees form a unique population.

Architectural analysis and deciphering growth strategies can also be applied to roots (Atger and Edelin 1994; Charles-Dominique et al. 2009). Root vs shoot allocation pattern can differ with the environment (Freschet et al. 2018). The root growth strategy directly drives to rooting depth, root lateral expansion, and root density (i.e. number of roots and root lengths per unit of soil volume), which will finally determine soil foraging capacity, water absorption capacity, and belowground intra- and interspecific competition.

Finally, our results were not in agreement with Borges et al. (2019), who applied a similar approach by comparing functional traits for an Asteraceae generalist tree species growing in savanna and cloud forests in a single site in south-eastern Brazil (the study was conducted on a same site, with no distance or climatic effects on functional traits). They found contrasting functional trait responses between the two habitats for a set of functional traits related to resource acquisition and storage (i.e. leaf area and thickness, LMA, wood density), such that savanna individuals were more resource conservative (i.e. high wood density and LMA, thick and small leaves) than those from cloud forest. The discrepancy between the two studies indicates that the type of phenotypic response (i.e. resource acquisition vs. resource use) for generalist species is not uniform across species, and may vary depending on its functional type (e.g. evergreen vs. deciduous, pioneer vs. late successional, light-demanding vs. shade tolerant), the nature of the resource heterogeneity between habitats (e.g. light, water, soil nutrients), or the range of the environmental gradient. Our study exemplifies the complexity of incorporating ITV in studying ecological processes, and how ITV of different functional traits are not evenly responsive to abiotic factors. However, we demonstrated the potential gains of incorporating architectural analysis in plant community ecology, particularly at the intraspecific level.

## Conclusion

Our study demonstrated that commonly measured traits, related to resource acquisition strategies, are not systematically responsive to contrasting habitats. Other aspects of plant functioning such as resource use strategies, such as through architectural development, can rather mediate such responses. Our study raises concerns about negative signal when investigating environmental filtering at the intraspecific level based on commonly measured functional traits like LMA, leaf thickness, or WSG. Environmental filtering can occur on other dimensions of plant functioning. As architectural analysis has brought insights on environmental filtering at the intraspecific level, such approach could also be applied to the process of niche differentiation, especially regarding intra- and interspecific competition.

## Data accessibility

Data are available online: https://zenodo.org/record/3632505#.XjQJqSPjK00

## Author contributions

PH designed and led the project. PH, SL, EN, VT, HM, NT, CS, SC, BF and HC measured tree architecture and functional traits. BF and VT described soils characteristics. SL, PH and GL performed data analysis. SL wrote the manuscript with contributions from PH and NT. All authors contributed critically to the drafts and gave final approval for publication.

### Acknowledgements

Version 4 of this preprint has been peer-reviewed and recommended by Peer Community In Ecology (https://doi.org/10.24072/pci.ecology.100041).

We especially thank Jean-Yves Goret, Saint-Omer Cazal and Audin Patient for their major contributions in field work and collecting data. We thank Julie Bossu, Coffi Belmys Cakpo, Jocelyn Cazal, Aurélie Cuvelier, Bruno Clair, Aurélie Dourdain, Alexandre Haslé de Barral, Solène Happert, Marie Hartwig, Clément Jouaux, Yohann Legraverant, Anabelle Mercrette, Ariane Mirabel, Pascal Petronelli, Laurent Risser, Dylan Taxile, Camille Ziegler and Lore Verryckt for their assistance with field work and measurement of leaf traits. We gratefully thank Isabelle Maréchaux and Christopher Baraloto for critical and valuable comments on the preliminary manuscript. We thank Anna Deasey for English proof reading. We thank the ONF for access to forestry roads and the sampling. We also thank the USRAVE (INRA-COFRAC) of Bordeaux and the SILVA lab (INRA) of Nancy for measurements of leaf nutrients.

S.L. was supported by a doctoral fellowship from CEBA. This study benefited from an Investissement d’Avenir grant managed by the Agence Nationale de la Recherche (CEBA, ref. ANR-10-LABX-0025).

## Conflict of interest disclosure

The authors of this preprint declare that they have no financial conflict of interest with the content of this article.

## Appendix

**Figure S1.**
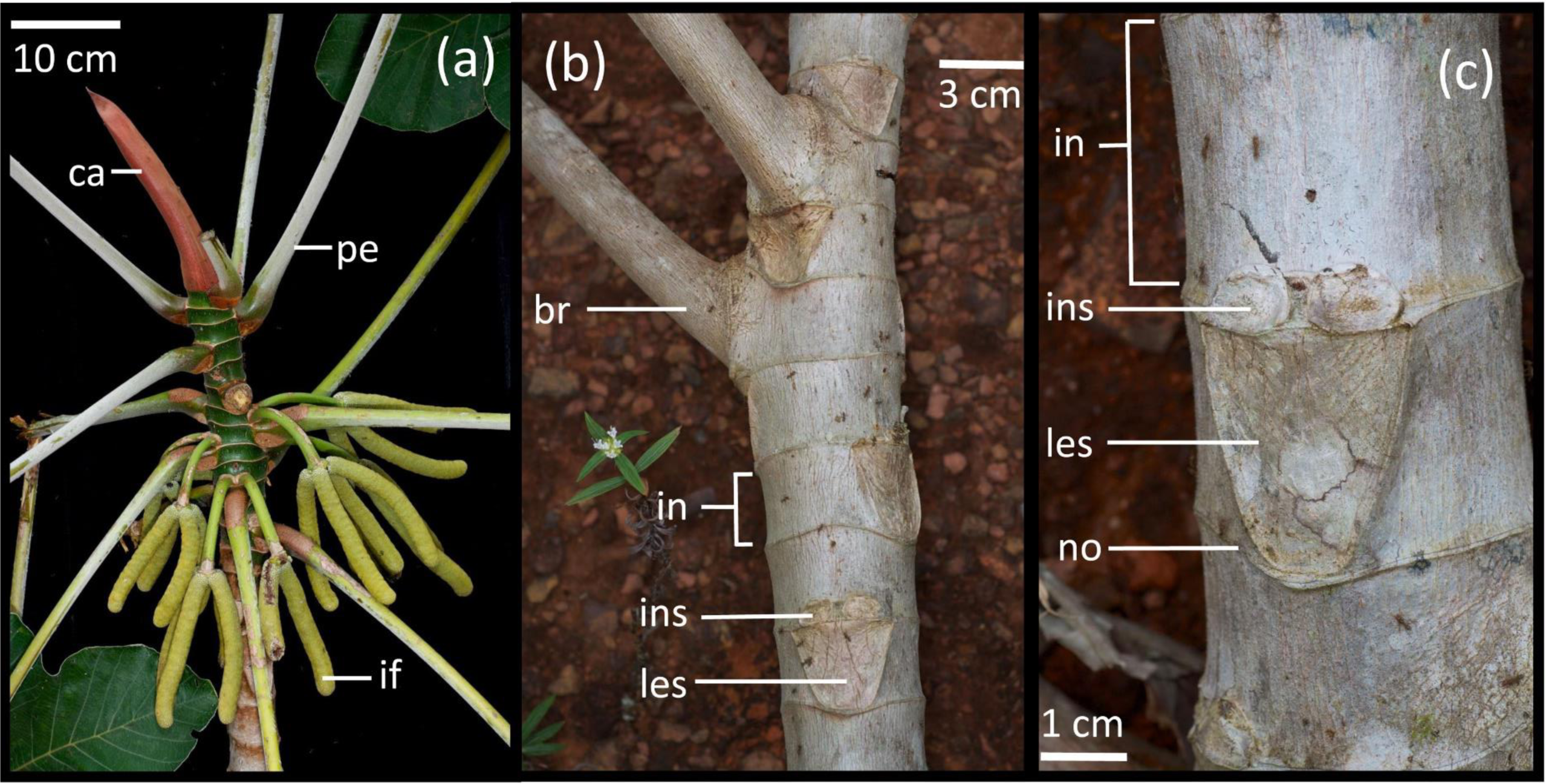
Morphological features of *Cecropia obtusa* Trécul (Urticaceae). (a) Focus on an apex, ca: calyptra; pe: petiole; if: inflorescence. (b) Focus on a branch tier, br: branch; in: internode, axis as the trunk are made of a linear succession of internodes; ins: inflorescence scars, these are twice just above the axillary leaf; les: leaf scar. (c) Focus on an internode, in: internode; ins: inflorescence scars; les: leaf scar; no: a node marled by the calyptra scar, allowing for the delineation of internodes along an axis as the trunk.

**Figure S2.**
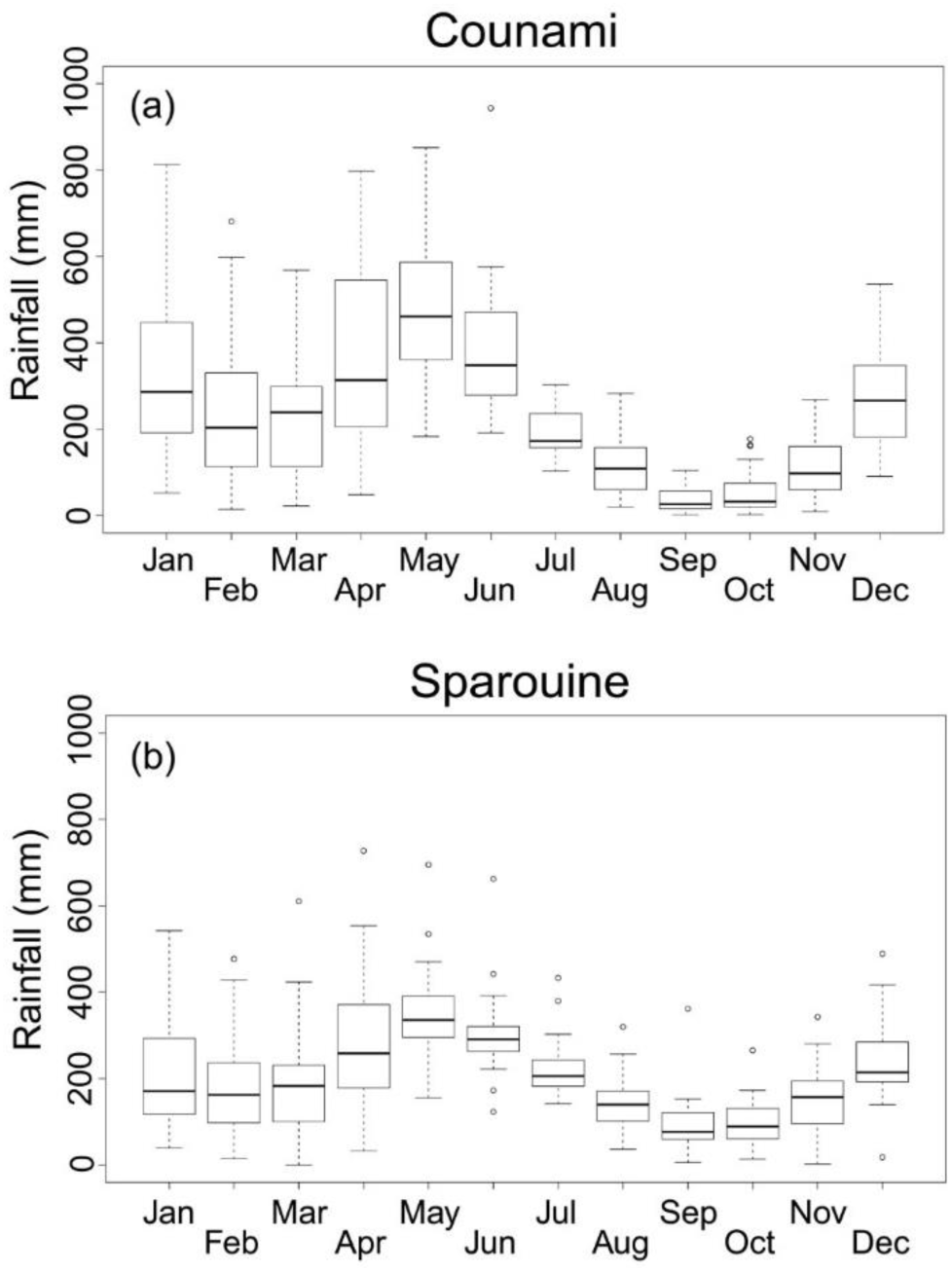
Box plots of mean annual rainfall (mm) from 1980 to 2016. (a) Counami, (b) Sparouine.

**Figure S3.**
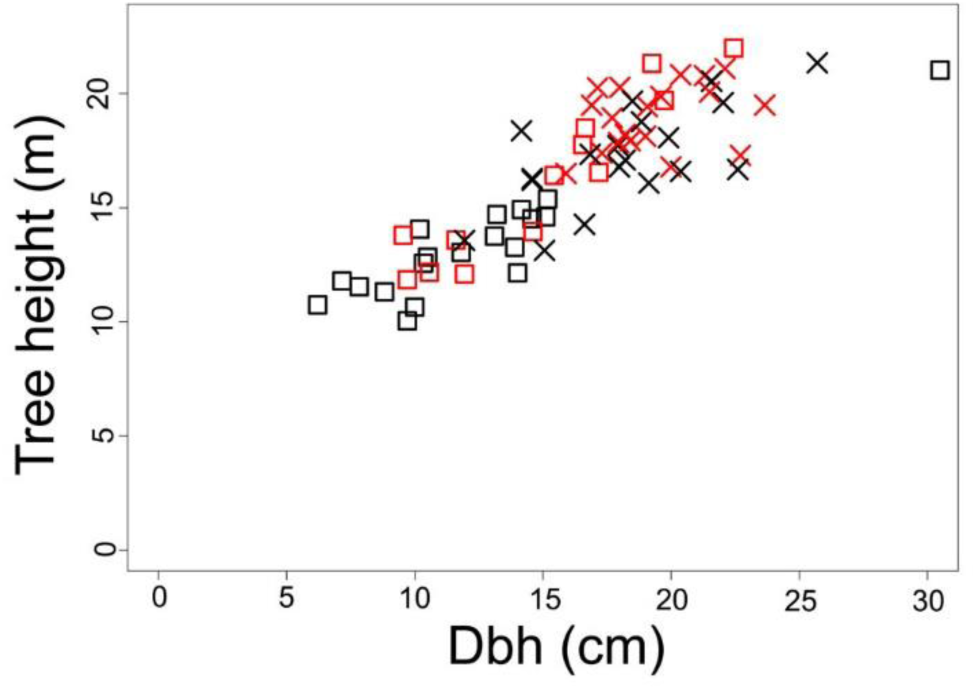
Tree heights (m) according to diameters at breast height (cm). Red: ferralitic soils; black: white-sand soils. Cross: Sparouine; square: Counami.

**Figure S4.**
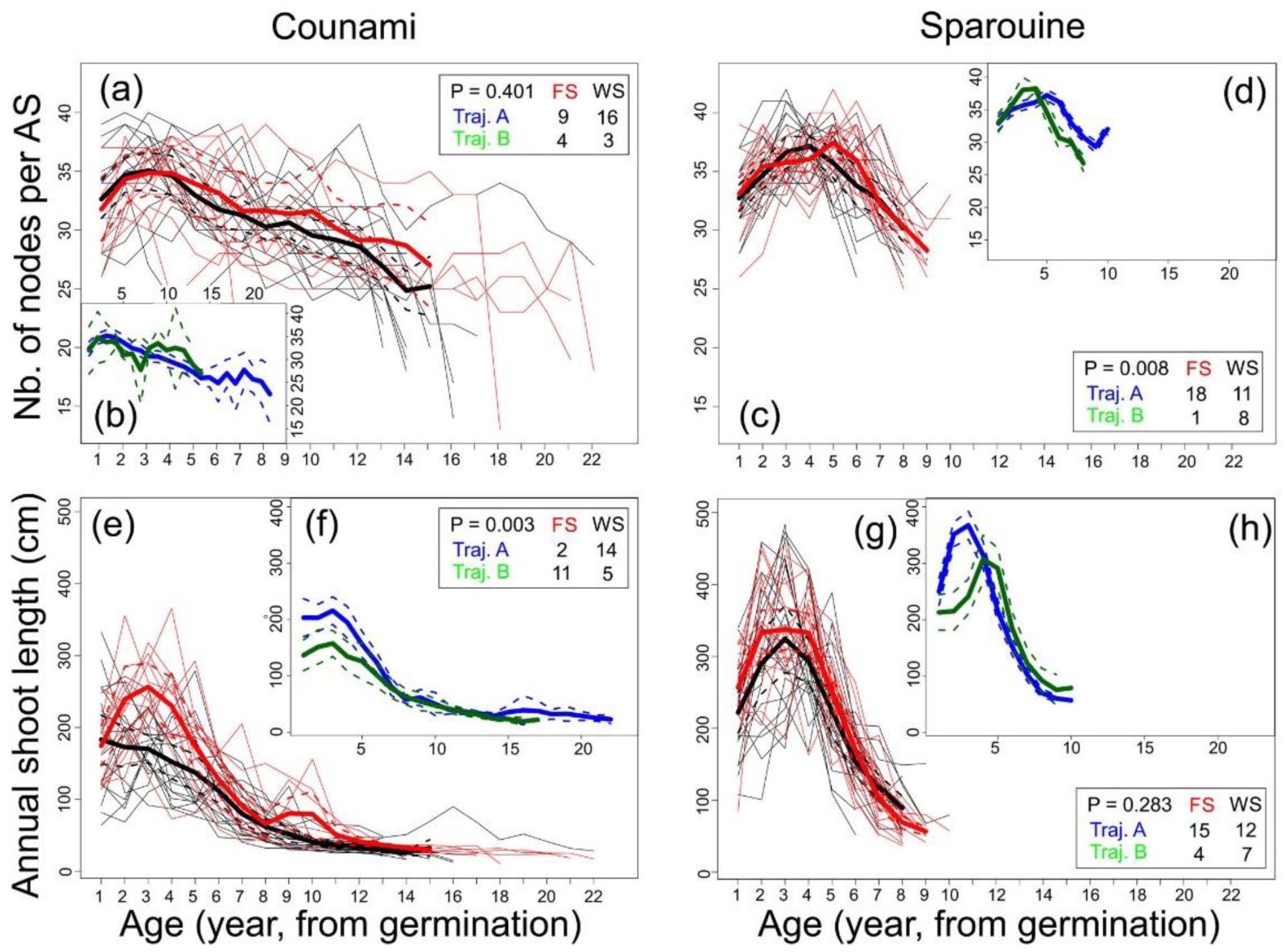
Other architectural growth traits according to age (years): number of nodes per annual shoot and annual shoot length. Main boxes represent features according to soil types. Inboxes represent mean trajectories after clustering longitudinal analyses (*kml*). The left column represents Counami trees, the right column represents Sparouine trees. Distributions between soil types and kml-trajectories are represented with Pearson chi-squared test. Red: ferralitic soils; black: white-sand soils. Blue: trajectory A; green: trajectory B; orange: trajectory C. Thick lines: means; dashed lines: confidence intervals at 95%.

### Appendix S1. Pedological characterization: Material and methods, and results

#### Material and methods

At Counami, soil sampling was conducted between the 8th and the 12th of September 2014, during the day. One sampling was conducted at the base of each individuals. At Sparouine, soil sampling was conducted between the 8th and the 10th of December 2014, during the day. Nine samples was harvested accounting for the 38 individuals because of the proximity of some trees (see Table S1.1). Per pool of trees, sampling was conducted for the individual at the most center of the considered pool. Sampling was conducted at the base of the considered individual, consisting in 4 excavation samples around the tree in cross, as there is a 2-m distance between two opposite excavated subsamples. Each subsamples are excavated at nearly 35 cm of depth. Nearly 500 g of soil material were sampled for each subsamples. For each tree, the 4 subsamples were mixed and dried in a closed room at 25°C and 50 % of relative humidity. Then, 50 g of sample were sent to LAS (Laboratoire d’Analyses des Sols), the central INRA (Institut National de la Recherche Agronomique) laboratory for soil analysis, at Arras for the measurement of soil properties presented in Table S1.2 with abbreviations.

**Table S1.1.**
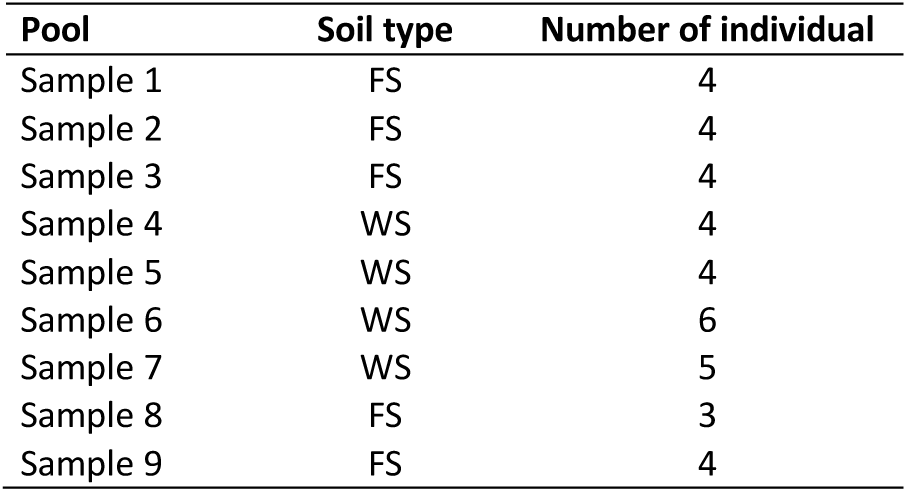
Soil sample pools for Sparouine

| Pool | Soil type | Number of individual |
| --- | --- | --- |
| Sample 1 | FS | 4 |
| Sample 2 | FS | 4 |
| Sample 3 | FS | 4 |
| Sample 4 | WS | 4 |
| Sample 5 | WS | 4 |
| Sample 6 | WS | 6 |
| Sample 7 | WS | 5 |
| Sample 8 | FS | 3 |
| Sample 9 | FS | 4 |

#### Results

The multivariate analysis of correlations of pedological properties defines two major axes (Fig. S1.1a) which explain 60.5 % of the inertia. The first axis (45.4 %) opposes (i) clayey and silty soils with high moisture, N, C, MO, P_tot_ contents and a high Al:CEC ratio to (ii) coarse sandy soils with high Ca:CEC and C:N ratios. The second axis (15.1 %) groups soils with respect to K:CEC, Fe:CEC, Mg:CEC, Na:CEC ratios and pH. When conditions are integrated (soil types x sites) the first axis reflects mainly the soil-type divergence (Fig. S1.1b), whereas the second one tends to reflect a between-site gradient. Within sites, the differences between soil types are more contrasted in Counami than in Sparouine. For a given soil type, the differences across sites are more contrasted between FS (P < 0.05, ANOVA, Fig. S1.1b) than between WS (P > 0.05, ANOVA, Fig. S1.1b), as the Sparouine FS are intermediary between Counami FS and all WS (P < 0.05, ANOVA, Fig. S1.1b). These features are reflected in ANOVA tests, where there are more significant differences in granulometric (clay, silt, sand) and organic matter (C, N, C:N, MO, P_tot_) properties between FS and WS (P < 0.05; Table S1.3), and between Sparouine FS and Counami FS (P < 0.05; Table S1.3), than Sparouine WS and Counami WS (P > 0.05; Table S1.3). Most of the exchangeable cations are only significantly differentiated according to soil types within the Counami site, or according to soil types and site at the same time. Otherwise, most of the exchangeable cations are not differentiated according to soil types within the Sparouine site.

**Table S1.2.** List of measured soil properties.

| Trait | Abbreviation | Unit |
| --- | --- | --- |
| *Water status* |  |  |
| Moisture content | H<sub>2</sub>O<sub>soil</sub> | g kg<sup>-1</sup> |
| *Granulometric* |  |  |
| Clay content | Clay | g kg<sup>-1</sup> |
| Thin silt content | Silt<sub>Th</sub> | g kg<sup>-1</sup> |
| Coarse silt content | Silt<sub>Co</sub> | g kg<sup>-1</sup> |
| Thin sand content | Sand<sub>Th</sub> | g kg<sup>-1</sup> |
| Coarse sand content | Sand<sub>Co</sub> | g kg<sup>-1</sup> |
| *Organic matter* |  |  |
| Carbon content | C<sub>soil</sub> | g kg<sup>-1</sup> |
| Nitrogen content | N<sub>soil</sub> | g kg<sup>-1</sup> |
| Carbon/Nitrogen ratio | CN<sub>soil</sub> | - |
| Organic matter content | MO<sub>soil</sub> | g kg<sup>-1</sup> |
| Total phosphorus content | P<sub>tot,soil</sub> | μg 100g<sup>-1</sup> |
| *Exchangeable cation* |  |  |
| Cation-exchangeable capacity | CEC | cmol kg<sup>-1</sup> |
| Calcium content | Ca<sub>soil</sub> | cmol kg<sup>-1</sup> |
| Magnesium content | Mg<sub>soil</sub> | cmol kg<sup>-1</sup> |
| Potassium content | K<sub>soil</sub> | cmol kg<sup>-1</sup> |
| Sodium content | Na<sub>soil</sub> | cmol kg<sup>-1</sup> |
| Aluminum content | Al<sub>soil</sub> | cmol kg<sup>-1</sup> |
| Hydrogen content | H<sub>soil</sub> | cmol kg<sup>-1</sup> |
| Iron content | Fe<sub>soil</sub> | cmol kg<sup>-1</sup> |
| Manganese content | Mn<sub>soil</sub> | cmol kg<sup>-1</sup> |
| pH | pH<sub>soil</sub> | - |

### Appendix S2. Material and methods: residuals, year delineation and inference of age

#### Residuals estimation

In the process of primary growth, three components are combined in an additive manner: an individual, an ontogenetic and an environmental component (Guédon et al. 2007, Chaubert-Pereira et al. 2009). To study the environmental component, based on the analysis of fluctuation of internode lengths, we used classical methods of time series analysis relying on a decomposition model principle. The ontogenetic (trend at low frequency) and environmental (fluctuations at mid frequency) sources of variation are differentiated by the application of a filter and then are separable. The applied linear filter converts measured sequences {*x*_*t*_} into another {*y*_*t*_} by a linear operation:

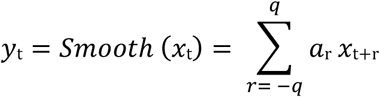

where {*a*_*r*_} is a set of weights such as for each *r, a*_*r*_ > 0 and *a*_*r*_ = *a*_*-r*_. To smooth out the local fluctuations and estimate the mean, weights should be chosen such that ∑_*r*_ *a*_*r*_ = 1. This linear operation is often referred to as a moving average, in which a smoothed curve is adjusted. To choose an appropriate filter is difficult and the choice remains partly arbitrary, so it is recommended to try a variety of filters to get an idea of the underlying trend. We chose to use the symmetric smoothing filter corresponding to the probability mass function of binomial distributions with parameters *n* = 200 and *p* = 0.05 to extract the trend of internode length sequences. Having extracting the trend, we looked at local fluctuations by examining the residuals. Residuals were extracted by division:

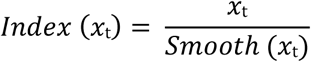

This standardization tends to allow us to give the same status to fluctuations of both small and large amplitudes, which is important in old trees that have very short internodes compared to the first node at the trunk base.

#### Year delineation and inference of age

For Sparouine trees, generated sequences of residuals are used for a cluster analysis with an alignment from the apex, since all Sparouine trees were felled at the same moment in the year. Resulting clusters gather trees for similar growth patterns and rainfall responses, and thus for similar growth rhythm. Generated mean trajectories of residual sequences are year-segmented based on the assumption that shorter internodes in one year are produced during the long dry season with lower rainfall, where the peak happens in September/October. Generated coordinated are directly injected for each individual according to its cluster belonging and adjusted if necessary.

For Counami trees, year delineation is directly realized on each individual’ sequence of residuals. Cluster analyses with an alignment from the apex are meaningless, because of contrasted felling dates over the year. Theoretically, year delineations for Counami trees are less robust.

For each year (between two year limits), knowing the number of nodes constituting this year, it is possible to estimate the mean number of days associated with the node production, namely the mean annual phyllochron, by dividing 365 by the number of constitutive nodes. In this way, we infer an inter-annual trend of phyllochron variation but no intra-annual variation. Finally, by cumulating the number of days associated with the production of each node, we estimate the age of each internode from germination. Then, it is possible to infer primary growth according to the chronological age. Within a year, the increase of age through successive internodes is regular and with the same interval.

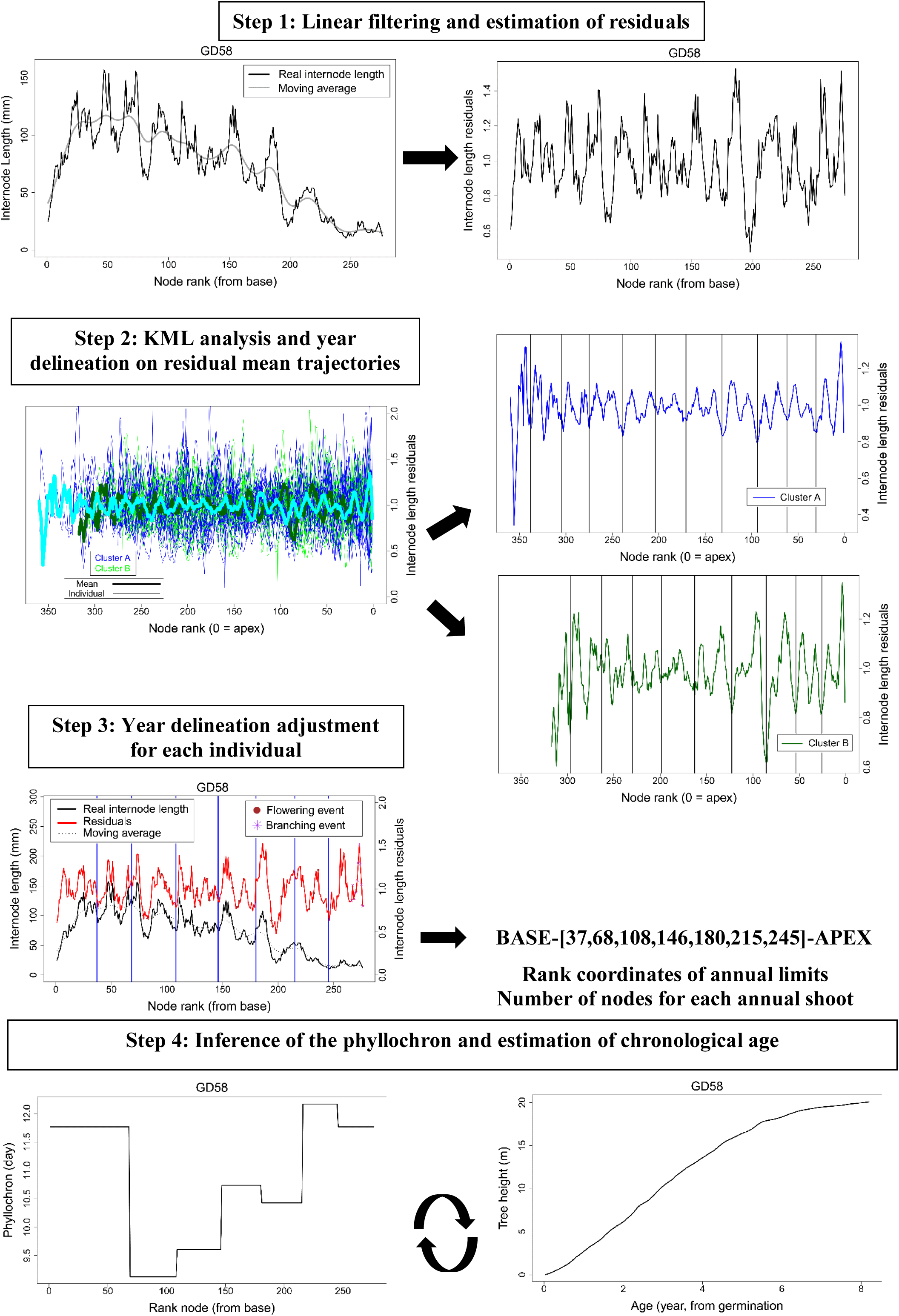

### Appendix S3. Materials and methods: measurement of leaf and trunk functional traits

Most of leaf functional traits (except those concerning leaf content traits) were measured the day when the tree was felled. The different steps are presented in the temporal order. The petiole length was measured on fresh leaves with a ruler. The petiole cross-sectional area was derived by measuring two orthogonal diameters from the fresh leaves at the exact middle of the petiole.

Leaf mass area (LMA) was measured by firstly cutting for each leaf, three tablets of known surface area (2.356 cm^2^), from the fresh limb at different points by avoiding veins. Secondly, tablets were dried at 70°C and the dry mass was measured. Finally, LMA was derived as follow:

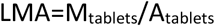

Leaf thickness was measured for each leaf on the fresh leaf at three different points by avoiding veins, with a micrometer. A mean is calculated with the three values. Since the *Cecropia obtusa* leaf is a palmatilobate leaf, lobes were counted. Leaf area was measured with a planimeter (LiCor 3000A, LiCor Inc, Lincoln, NE, USA). Foliar chlorophyll content was assessed with a SPAD-502 chlorophyll meter (Konica Minolta, Osaka, Japan) with a specific *Cecropia obtusa* calibration (Coste et al. 2010). Wood specific gravity (WSG) is estimated with a complete wood slice sampled at 1 meter of height. Bark is removed and wood specific gravity is estimated with the method described by (Williamson and Wiemann 2010):

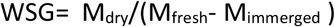

Where M_dry_ is the wood slice dry mass (g), M_fresh_ is the wood slice fresh mass (g), and M_immerged_ is the wood slice mass (g) with the slice put under water. M_fresh_ was measured the day when the tree was felled. M_dry_ was measured after the slice was dried at 103°C.

Leaf lifespan was deviated on the postulate of a constant-10-day phyllochron (i.e. rhythm of leaf production) as shown by Heuret et al. (2002) by multiplying the number of leaves of the A1 axis by 10 days.

Since most of leaves were damaged causing an underestimation of leaf area measurements, we rely on a prediction model to calculate an estimated leaf area and thus an estimated crown area, since the relationship between the length of the main lobe and the undamaged lamina area is very informative (Levionnois et al., 2020; R^2^ = 0.942; P < 0.001). The model is based on 1064 mature and undamaged leaves and shaped from a NLS (non-linear least squares) with the basic STATS package performed with R software (http://CRAN-R-project.org). For each tree of the study, the length of the main lobe was measured for every leaf of the crown with a ruler. Based on our model, the crown area was estimated as the sum of every estimated leaf area.

For leaf content traits, leaves were dried at 70°C. Nearly 10g of dry matters are sampled for each leaf, by avoiding veins. Samples were sent to USRAVE (Unité de Service et de Recherche en Analyses Végétales et Environnementales), the central INRA (Institut National de la Recherche Agronomique) laboratory for plant analysis accredited by COFRAC (Comité Français d’Accréditation), at Bordeaux. Measurements of leaf content traits were conducted for the residual moisture content, 13 ⍰C, carbon, nitrogen, phosphorus and potassium.

### Appendix S4. Autocorrelation coefficients: methods and results

#### Material and methods

We relied on sample autocorrelation coefficients to point out a potential annual periodicity on the stand level (i.e. soil x site) for growth, branching, and flowering process. Thereafter, such stand periodicity would help to improve retrospective analysis of tree development with a temporal scale. The use of sample autocorrelation coefficients for all trees together allows measurement of the correlation between observations of sequences of quantitative variables separated by different distances. The autocorrelation function measures the correlation between *X*_*t*_ and *X*_*t+k*_ as a function of the internode lag k. The sample autocorrelation function is an even function of the internode lag and hence needs to be plotted for *k* = 0, 1, 2, …, *n*. We applied auto-correlation analysis to residual sequences obtained from internode length sequences after removing the ontogenetic trend, to binary branching, and flowering sequences (Guédon et al. 2001, 2003).

#### Results

Autocorrelation functions calculated for the internode length residues as well as branching and flowering binary sequences all showed significant periodicity, independently of the site or soil type (Fig. S4.1). In Counami, the correlogram calculated for internode residues sequences yielded significant positive maxima at lags 30, 58, 88 in WS and 30, 52, 64, 88 in FS (Fig. S4.1). In Sparouine, a similar pattern was observed with significant positive maxima at lags 17, 34, 79 in WS and 19, 36, 68, 88 in FS (Fig. S4.1b). Similar overall patterns are observed for flowering event sequences. In Counami, the correlogram yielded significant positive maxima at lags 12, 27, 52, 77, 88 in WS and 27, 53, 73, 89 in FS (Fig. S4.1c). In Sparouine, significant positive maxima were at lags 16, 32, 63, 74 in WS and 19, 33, 48, 64, 89 in FS (Fig. S4.1d). Considering the first 50 lags, a bimodal pattern (lags 12-19 and 27-36) is more pronounced in Sparouine and on WS for internode length and flowering variables. In Counami, the correlogram calculated for branching event sequences yielded significant positive maxima at lags 38, 60 in WS and 29, 60, 88 in FS (Fig. S4.1e). In Sparouine, the correlogram calculated for this variable yielded significant positive maxima at lags 30, 40, 70 in WS and 28, 38, 100 in FS (Fig. S4.1f).

Based on rhythms relying on autocorrelation coefficients and the knowledge on *C. obtusa* (Heuret et al. 2002) and *C. sciadophylla* (Zalamea et al. 2008), we delineated years, trying to respect the following rules: 30-35 trunk nodes per year, 1 to 2 trunk flowering events per year, and 0 or 1 trunk branching event (i.e. tiers of branches) per year. Based on such diagnosis, we calculated the date of the formation of each node and switched from a topological scale, the rank of the node, to a temporal scale (Appendix S2).

---

**Fig. S4.1.**
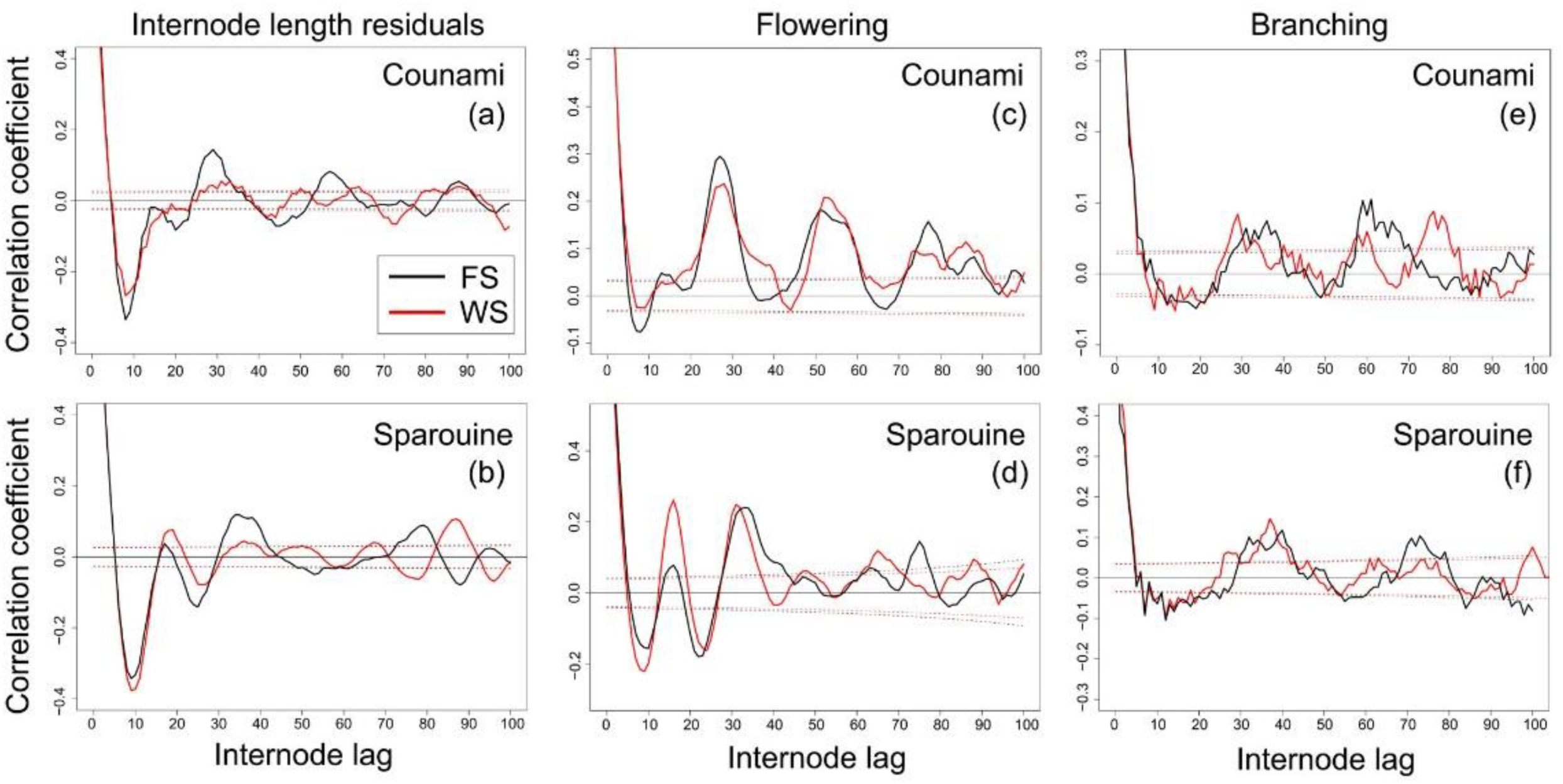
Autocorrelation function according to growth, flowering, and branching processes. (a) Internode length residuals for Counami, (b) Internode length residuals for Sparouine, (c) Flowering presence for Counami, (d) Flowering presence for Sparouine, (e) Branching presence for Counami, (f) Branching presence for Sparouine. Red: ferralitic soils; black: white-sand soils.

## Notes

#### Summary of Updates

Materials and Methods, and Discussion have been clarified.

https://zenodo.org/record/3632505#.XjQJqSPjK00

## References

Adams HD, Zeppel MJB, Anderegg WRL, et al. 2017. A multi-species synthesis of physiological mechanisms in drought-induced tree mortality. Nature Ecology and Evolution.

Adeney JM, Christensen NL, Vicentini A, Cohn-Haft M. 2016. White-sand Ecosystems in Amazonia. Biotropica 48: 7–23.

Allié E, Pélissier R, Engel J, et al. 2015. Pervasive Local-Scale Tree-Soil Habitat Association in a Tropical Forest Community. PLOS ONE 10: e0141488.

Anderegg WRL, Klein T, Bartlett M, et al. 2016. Meta-analysis reveals that hydraulic traits explain cross-species patterns of drought-induced tree mortality across the globe. Proceedings of the National Academy of Sciences of the United States of America 113: 5024–5029.

Atger C, Edelin C. 1994. Premières données sur l’architecture comparée des systèmes racinaires et caulinaires. Canadian Journal of Botany 72: 963–975.

Baraloto C, Timothy Paine CE, Poorter L, et al. 2010. Decoupled leaf and stem economics in rain forest trees. Ecology Letters 13: 1338–1347.

Bartlett MK, Scoffoni C, Sack L. 2012. The determinants of leaf turgor loss point and prediction of drought tolerance of species and biomes: a global meta-analysis. Ecology Letters 15: 393–405.

Bettiati D, Petit G, Anfodillo T. 2012. Testing the equi-resistance principle of the xylem transport system in a small ash tree: empirical support from anatomical analyses. Tree Physiology 32: 171–177.

Borges ER, Prado-Junior J, Santana LD, et al. 2019. Trait variation of a generalist tree species (Eremanthus erythropappus, Asteraceae) in two adjacent mountain habitats: savanna and cloud forest. Australian Journal of Botany 66: 640–646.

Borregaard MK, Rahbek C. 2010. Causality of the relationship between geographic distribution and species abundance. The Quarterly Review of Biology 85: 3–25.

Boulangeat I, Lavergne S, Es JV, Garraud L, Thuiller W. 2012. Niche breadth, rarity and ecological characteristics within a regional flora spanning large environmental gradients. Journal of Biogeography 39: 204–214.

Charles-Dominique T, Mangenet T, Rey H, Jourdan C, Edelin C. 2009. Architectural analysis of root system of sexually vs. vegetatively propagated yam (Dioscorea rotundata Poir.), a tuber monocot. Plant and Soil 317: 61–77.

Chessel D, Dufour A-B, Thioulouse J. 2004. The ade4 package - I: One-table methods. R News 4: 5–10.

Clark DB, Palmer MW, Clark DA. 1999. Edaphic Factors and the Landscape-Scale Distributions of Tropical Rain Forest Trees. Ecology 80: 2662–2675.

Coste S, Roggy J-C, Garraud L, Heuret P, Nicolini E, Dreyer E. 2009. Does ontogeny modulate irradiance-elicited plasticity of leaf traits in saplings of rain-forest tree species? A test with Dicorynia guianensis and Tachigali melinonii (Fabaceae, Caesalpinioideae). Annals of Forest Science 66: 709–709.

Daly DC, Silveira M, Medeiros H, Castro W, Obermüller FA. 2016. The White-sand Vegetation of Acre, Brazil. Biotropica 48: 81–89.

Dang-Le AT, Edelin C, Le-Cong K. 2013. Ontogenetic variations in leaf morphology of the tropical rain forest species Dipterocarpus alatus Roxb. ex G. Don. Trees 27: 773–786.

Davis RB. 1970. Seasonal differences in intermodal lengths in Cecropia trees; a suggested method for measurement of past growth in height. Turrialba.

Dejean A, Grangier J, Leroy C, Orivel J. 2009. Predation and aggressiveness in host plant protection: a generalization using ants from the genus Azteca. Naturwissenschaften 96: 57–63.

Eller C, de V. Barros F, R.L. Bittencourt P, Rowland L, Mencuccini M, S. Oliveira R. 2018. Xylem hydraulic safety and construction costs determine tropical tree growth. Plant, Cell & Environment: n/a-n/a.

Fine PVA, Baraloto C. 2016. Habitat Endemism in White-sand Forests: Insights into the Mechanisms of Lineage Diversification and Community Assembly of the Neotropical Flora. Biotropica 48: 24–33.

Fine PVA, García-Villacorta R, Pitman NCA, Mesones I, Kembel SW. 2010. A floristic study of the white-sand forests of Peru. Annals of the Missouri Botanical Garden 97: 283–305.

Fine PVA, Mesones I, Coley PD. 2004. Herbivores Promote Habitat Specialization by Trees in Amazonian Forests. Science 305: 663–665.

Fine PVA, Metz MR, Lokvam J, et al. 2013. Insect herbivores, chemical innovation, and the evolution of habitat specialization in Amazonian trees. Ecology 94: 1764–1775.

Fine PVA, Miller ZJ, Mesones I, et al. 2006. The growth–defense trade-off and habitat specialization by plants in amazonian forests. Ecology 87: S150–S162.

Folgarait PJ, Davidson DW. 1994. Antiherbivore defenses of myrmecophytic Cecropia under different light regimes. Oikos 71: 305–320.

Folgarait PJ, Davidson DW. 1995. Myrmecophytic Cecropia: antiherbivore defenses under different nutrient treatments. Oecologia 104: 189–206.

Fortunel C, Fine PVA, Baraloto C. 2012. Leaf, stem and root tissue strategies across 758 Neotropical tree species. Functional Ecology 26: 1153–1161.

Fortunel C, Paine CET, Fine PVA, Kraft NJB, Baraloto C. 2014. Environmental factors predict community functional composition in Amazonian forests. Journal of Ecology 102: 145–155.

Fortunel C, Ruelle J, Beauchêne J, Fine PVA, Baraloto C. 2014. Wood specific gravity and anatomy of branches and roots in 113 Amazonian rainforest tree species across environmental gradients. New Phytologist 202: 79–94.

Freschet GT, Valverde-Barrantes OJ, Tucker CM, et al. 2017. Climate, soil and plant functional types as drivers of global fine-root trait variation. Journal of Ecology 105: 1182–1196.

Freschet GT, Violle C, Bourget MY, Scherer-Lorenzen M, Fort F. 2018. Allocation, morphology, physiology, architecture: the multiple facets of plant above- and below-ground responses to resource stress. New Phytologist 219: 1338–1352.

Fridley JD, Grime JP. 2010. Community and ecosystem effects of intraspecific genetic diversity in grassland microcosms of varying species diversity. Ecology 91: 2272–2283.

Fyllas NM, Patiño S, Baker TR, et al. 2009. Basin-wide variations in foliar properties of Amazonian forest: phylogeny, soils and climate. Biogeosciences 6: 2677–2708.

Genolini C, Falissard B. 2009. KmL: k-means for longitudinal data. Computational Statistics 25: 317–328.

Godin C, Caraglio Y. 1998. A Multiscale Model of Plant Topological Structures. Journal of Theoretical Biology 191: 1–46.

Godin C, Costes E, Caraglio Y. 1997. Exploring plant topological structure with the AMAPmod software: an outline.

Gourlet-Fleury S, Guehl JM, Laroussine O. 2004. Ecology and management of a neotropical rainforest: lessons drawn from Paracou, a long-term experimental research site in French Guiana. Paris: Elsevier.

Grubb PJ, Coomes DA. 1997. Seed mass and nutrient content in nutrient-starved tropical rainforest in Venezuela. Seed Science Research 7: 269–280.

Guédon Y, Caraglio Y, Heuret P, Lebarbier E, Meredieu C. 2007. Analyzing growth components in trees. Journal of Theoretical Biology 248: 418–447.

Heuret P, Barthélémy D, Guédon Y, Coulmier X, Tancre J. 2002. Synchronization of growth, branching and flowering processes in the South American tropical tree Cecropia obtusa (Cecropiaceae). American Journal of Botany 89: 1180–1187.

Heuret P, Meredieu C, Coudurier T, Courdier F, Barthélémy D. 2006. Ontogenetic trends in the morphological features of main stem annual shoots of Pinus pinaster (Pinaceae). American Journal of Botany 93: 1577–1587.

HilleRisLambers J, Adler PB, Harpole WS, Levine JM, Mayfield MM. 2012. Rethinking Community Assembly through the Lens of Coexistence Theory. Annual Review of Ecology, Evolution, and Systematics 43: 227–248.

Holt AR, Gaston KJ, He F. 2002. Occupancy-abundance relationships and spatial distribution: A review. Basic and Applied Ecology 3: 1–13.

Jung V, Albert CH, Violle C, Kunstler G, Loucougaray G, Spiegelberger T. 2014. Intraspecific trait variability mediates the response of subalpine grassland communities to extreme drought events. Journal of Ecology 102: 45–53.

Jung V, Violle C, Mondy C, Hoffmann L, Muller S. 2010. Intraspecific variability and trait-based community assembly. Journal of Ecology 98: 1134–1140.

Kassambara A, Mundt F. 2016. Factoextra: Extract and Visualize the Results of Multivariate Data Analyses.

Kraft NJB, Adler PB, Godoy O, James EC, Fuller S, Levine JM. 2015. Community assembly, coexistence and the environmental filtering metaphor. Functional Ecology 29: 592–599.

Kraft NJB, Crutsinger GM, Forrestel EJ, Emery NC. 2014. Functional trait differences and the outcome of community assembly: an experimental test with vernal pool annual plants. Oikos 123: 1391–1399.

Kraft NJB, Valencia R, Ackerly DD. 2008. Functional Traits and Niche-Based Tree Community Assembly in an Amazonian Forest. Science 322: 580–582.

Latteman TA, Mead JE, DuVall MA, Bunting CC, Bevington JM. 2014. Differences in anti-herbivore defenses in non-myrmecophyte and myrmecophyte Cecropia trees. Biotropica 46: 652–656.

Lehnebach R, Bossu J, Va S, et al. 2019. Wood Density Variations of Legume Trees in French Guiana along the Shade Tolerance Continuum: Heartwood Effects on Radial Patterns and Gradients. Forests 10: 80.

Lepš J, Bello F de, Šmilauer P, Doležal J. 2011. Community trait response to environment: disentangling species turnover vs intraspecific trait variability effects. Ecography 34: 856–863.

Letort V, Heuret P, Zalamea P-C, Reffye PD, Nicolini E. 2012. Analysing the effects of local environment on the source-sink balance of Cecropia sciadophylla: a methodological approach based on model inversion. Annals of Forest Science 69: 167–180.

Marschner H. 1995. 8 - Functions of Mineral Nutrients: Macronutrients In: Mineral Nutrition of Higher Plants (Second Edition). London: Academic Press, 229–312.

Mathieu A, Letort V, Cournède P h., Zhang B g., Heuret P, de Reffye P. 2012. Oscillations in Functional Structural Plant Growth Models. Mathematical Modelling of Natural Phenomena 7: 47–66.

McGill BJ, Enquist BJ, Weiher E, Westoby M. 2006. Rebuilding community ecology from functional traits. Trends in Ecology & Evolution 21: 178–185.

Niklas KJ. 2007. Maximum plant height and the biophysical factors that limit it. Tree Physiology 27: 433–440.

O’Brien MJ, Engelbrecht BMJ, Joswig J, et al. 2017. A synthesis of tree functional traits related to drought-induced mortality in forests across climatic zones. Journal of Applied Ecology.

Oldham AR, Sillett SC, Tomescu AMF, Koch GW. 2010. The hydrostatic gradient, not light availability, drives height-related variation in Sequoia sempervirens (Cupressaceae) leaf anatomy. American Journal of Botany 97: 1087–1097.

Paine CET, Baraloto C, Chave J, Hérault B. 2011. Functional traits of individual trees reveal ecological constraints on community assembly in tropical rain forests. Oikos 120: 720–727.

Patiño S, Lloyd J, Paiva R, et al. 2009. Branch xylem density variations across the Amazon Basin. Biogeosciences 6: 545–568.

Pradal C, Coste J, Boudon F, Fournier C, Godin C. 2013. OpenAlea 2.0: Architecture of an integrated modeling environment on the web. Finnish Society of Forest Science.

Prendin AL, Mayr S, Beikircher B, von Arx G, Petit G. 2018. Xylem anatomical adjustments prioritize hydraulic efficiency over safety as Norway spruce trees grow taller. Tree Physiology 38: 1088–1097.

R Core Team. 2018. R: A language and environment for statistical computing. Vienna, Austria: R Foundation for Statistical Computing.

Reich PB. 2014. The world-wide ‘fast–slow’ plant economics spectrum: a traits manifesto. Journal of Ecology 102: 275–301.

Roggy J-C, Nicolini É, Imbert P, Caraglio Y, Bosc A, Heuret P. 2005. Links between tree structure and functional leaf traits in the tropical forest tree Dicorynia guianensis Amshoff (Caesalpiniaceae).

Roy M, Schimann H, Braga-Neto R, et al. 2016. Diversity and Distribution of Ectomycorrhizal Fungi from Amazonian Lowland White-sand Forests in Brazil and French Guiana. Biotropica 48: 90–100.

Rungwattana Kanin, Hietz Peter, Larjavaara Markku. 2017. Radial variation of wood functional traits reflect size-related adaptations of tree mechanics and hydraulics. Functional Ecology 32: 260–272.

Ryan MG, Phillips N, Bond BJ. 2006. The hydraulic limitation hypothesis revisited. Plant, Cell & Environment 29: 367–381.

Sabatier D, Grimaldi M, Prévost M-F, et al. 1997. The influence of soil cover organization on the floristic and structural heterogeneity of a Guianan rain forest. Plant Ecology 131: 81–108.

Schamp BS, Chau J, Aarssen LW. 2008. Dispersion of traits related to competitive ability in an old-field plant community. Journal of Ecology 96: 204–212.

Schupp EW. 1986. Azteca protection of Cecropia: ant occupation benefits juvenile trees. Oecologia 70: 379–385.

Shipley B, Bello FD, Cornelissen JHC, Laliberté E, Laughlin DC, Reich PB. 2016. Reinforcing loose foundation stones in trait-based plant ecology. Oecologia 180: 923–931.

Sides CB, Enquist BJ, Ebersole JJ, Smith MN, Henderson AN, Sloat LL. 2014. Revisiting Darwin’s hypothesis: Does greater intraspecific variability increase species’ ecological breadth? American Journal of Botany 101: 56–62.

ter Steege H, Pitman NCA, Sabatier D, et al. 2013. Hyperdominance in the Amazonian Tree Flora. Science 342: 1243092.

Stropp J, Sleen PV der, Assunção PA, Silva AL da, Steege HT. 2011. Tree communities of white-sand and terra-firme forests of the upper Rio Negro. Acta Amazonica 41: 521–544.

Swenson NG, Enquist BJ. 2009. Opposing assembly mechanisms in a Neotropical dry forest: implications for phylogenetic and functional community ecology. Ecology 90: 2161–2170.

Taugourdeau O, Dauzat J, Griffon S, Sabatier S, Caraglio Y, Barthélémy D. 2012. Retrospective analysis of tree architecture in silver fir (Abies alba Mill.): ontogenetic trends and responses to environmental variability. Annals of Forest Science 69: 713–721.

Uriarte M, Condit R, Canham CD, Hubbell SP. 2004. A spatially explicit model of sapling growth in a tropical forest: does the identity of neighbours matter? Journal of Ecology 92: 348–360.

Urli M, Porté AJ, Cochard H, Guengant Y, Burlett R, Delzon S. 2013. Xylem embolism threshold for catastrophic hydraulic failure in angiosperm trees. Tree Physiology 33: 672–683.

Violle C, Enquist BJ, McGill BJ, et al. 2012. The return of the variance: intraspecific variability in community ecology. Trends in Ecology & Evolution 27: 244–252.

Violle C, Navas M-L, Vile D, et al. 2007. Let the concept of trait be functional! Oikos 116: 882–892.

Wagner F, Rossi V, Stahl C, Bonal D, Hérault B. 2012. Water Availability Is the Main Climate Driver of Neotropical Tree Growth. PLOS ONE 7: e34074.

Zalamea P-C, Heuret P, Sarmiento C, et al. 2012. The Genus Cecropia: A Biological Clock to Estimate the Age of Recently Disturbed Areas in the Neotropics. PLoS ONE 7: e42643.

Zalamea P-C, Sarmiento C, Stevenson PR, Rodríguez M, Nicolini E, Heuret P. 2013. Effect of rainfall seasonality on the growth of Cecropia sciadophylla: intra-annual variation in leaf production and node length. Journal of Tropical Ecology 29: 361–365.

Zalamea P-C, Stevenson PR, Madriñán S, Aubert P-M, Heuret P. 2008. Growth pattern and age determination for Cecropia sciadophylla (Urticaceae). American Journal of Botany 95: 263–271.

## Literature cited

Chaubert-Pereira, F., Y. Caraglio, C. Lavergne, and Y. Guédon. 2009. Identifying ontogenetic, environmental and individual components of forest tree growth. Annals of Botany 104:883–896.

Guédon, Y., Y. Caraglio, P. Heuret, E. Lebarbier, and C. Meredieu. 2007. Analyzing growth components in trees. Journal of Theoretical Biology 248:418–447.

## Literature cited

Coste, S., C. Baraloto, C. Leroy, É. Marcon, A. Renaud, A. D. Richardson, J.-C. Roggy, H. Schimann, J. Uddling, and B. Hérault. 2010. Assessing foliar chlorophyll contents with the SPAD-502 chlorophyll meter: a calibration test with thirteen tree species of tropical rainforest in French Guiana. Annals of Forest Science 67:607–607.

Heuret, P., D. Barthélémy, Y. Guédon, X. Coulmier, and J. Tancre. 2002. Synchronization of growth, branching and flowering processes in the South American tropical tree Cecropia obtusa (Cecropiaceae). American Journal of Botany 89:1180–1187.

Levionnois, S., S. Coste, E. Nicolini, C. Stahl, H. Morel, and P. Heuret. Scaling of petiole anatomies, mechanics, and vasculatures with leaf size in the widespread Neotropical pioneer tree species Cecropia obtusa Trécul (Urticaceae), Tree Physiology, tpz136

Williamson, G. B., and M. C. Wiemann. 2010. Measuring wood specific gravity…Correctly. American Journal of Botany 97:519–524.

## Literature cited

Guédon Y, Barthélémy D, Caraglio Y, Costes E. 2001. Pattern analysis in branching and axillary flowering sequences. Journal of Theoretical Biology 212: 481–520.

Guédon Y, Heuret P, Costes E. 2003. Comparison methods for branching and axillary flowering sequences. Journal of Theoretical Biology 225: 301–325.

